# HMGB1 restores a dynamic chromatin environment in the presence of linker histone by deforming nucleosomal DNA

**DOI:** 10.1101/2024.08.23.609244

**Authors:** Hayden S. Saunders, Un Seng Chio, Camille M. Moore, Vijay Ramani, Yifan Cheng, Geeta J Narlikar

**Author notes:** These authors contributed equally (Un Seng Chio, Camille M. Moore).

## Abstract

The essential architectural protein HMGB1 increases accessibility of nucleosomal DNA and counteracts the effects of linker histone H1. However, HMGB1 is less abundant than H1 and binds nucleosomes more weakly raising the question of how HMGB1 effectively competes with H1. Here, we show that HMGB1 rescues H1’s inhibition of nucleosomal DNA accessibility without displacing H1. HMGB1 also increases the dynamics of condensed, H1-bound chromatin. Cryo-EM shows that HMGB1 binds at internal locations on a nucleosome and locally distorts the DNA. These sites, which are away from the binding site of H1, explain how HMGB1 and H1 co-occupy a nucleosome. Our findings lead to a model where HMGB1 counteracts the activity of H1 by distorting nucleosomal DNA and by contacting the H1 C-terminal tail. Compared to direct competition, nucleosome co-occupancy by HMGB1 and H1 allows a greater diversity of dynamic chromatin states and may be generalizable to other chromatin regulators.

## Introduction

In eukaryotes, the fundamental unit of chromatin is a nucleosome, which is composed of ∼147 base pairs (bp) of DNA wrapped around an octamer of histone proteins. Processes such as transcription, replication or DNA repair that require access to DNA often compete with the histone octamer (Lorch et al., 1987). It is well known that ATP-dependent chromatin remodelers and histone chaperones promote access to nucleosomal DNA by sliding or disassembling nucleosomes to stably expose tracts of DNA (Clapier et al., 2017; Winkler and Luger, 2011). However, the intrinsic dynamics of chromatin also regulate global DNA accessibility (Anderson and Widom, 2000; Armeev et al., 2021; Bilokapic et al., 2018a, 2018b; Polach and Widom, 1995). In this context, various nuclear architectural proteins that modify chromatin dynamics are implicated in regulating a wide variety of nuclear processes that need access to DNA (Fyodorov et al., 2018; Thomas and Stott, 2012; Thomas and Travers, 2001). One such architectural protein, high mobility group box 1 (HMGB1), is highly abundant in the eukaryotic nucleus (Duguet and de Recondo, 1978). *In vivo* HMGB1 is associated with chromatin decompaction and transcriptional activation (Aizawa et al., 1994; Ogawa et al., 1995). These activities are consistent with HMGB1 granting access to nucleosomal DNA and are thought to be opposed by another abundant nuclear architectural protein, linker histone H1, which is associated with chromatin compaction and transcriptional repression (Fyodorov et al., 2018; Ragab and Travers, 2003). Yet how these activities compete at the nucleosome level and how they translate to effects at the scale of chromatin remains unclear.

It is proposed that HMGB1 and H1 share an overlapping binding site on the nucleosome near the entry/exit site of DNA such that competition for binding regulates their relative activities (An et al., 1998; Nightingale et al., 1996; Cato et al., 2008; Catez et al., 2004; Thomas and Stott, 2012). Indeed, examples of transitions from HMGB1-bound chromatin to H1-bound chromatin or vice versa have been observed during *Xenopus* and *Drosophila* embryonic development and at gene promoters during transcription initiation (Ju et al., 2006; Ner and Travers, 1994; Nightingale et al., 1996). Additionally, H1 turnover dynamics are sensitive to HMGB1 concentration *in vivo,* suggestive of competition through overlapping binding sites (Catez et al., 2004). However, HMGB1 is present in the nucleus at 10-fold lower concentrations and has ∼1000-fold weaker affinity for nucleosomes and shorter residence times on chromatin than H1 (Bonaldi et al., 2002; Duguet and de Recondo, 1978; Falciola et al., 1997; Phair et al., 2004; Scaffidi et al., 2002; Ueda et al., 2004; White et al., 2016). These differences raise the question of how HMGB1 effectively competes for the same binding site with H1. The binding competition model is further complicated by the observation that HMGB1 and H1 interact with one another (Cato et al., 2008). Direct competition between HMGB1 and H1 has not been studied *in vitro* on nucleosomes, so other modes of competition have not been tested. HMGB1’s attributes on nucleosomes are likely due to its domain structure and unique mode of DNA binding. HMGB1 consists of two HMG box domains, which bind DNA without sequence specificity. Each box contains two hydrophobic residues that intercalate into the minor groove of DNA and induce a bend in the DNA (Thomas and Travers, 2001). HMGB1 also contains a C-terminal autoinhibitory domain consisting of 30 consecutive negatively charged Asp and Glu residues, which mimics DNA to occlude the DNA binding interface of the HMG boxes (Figure 1A) (Knapp et al., 2004; Stott et al., 2010; Watson et al., 2007). Therefore, DNA bound by HMGB1 is in competition with the auto-inhibitory C-terminal tail, resulting in a high off rate and low affinity. These binding and bending dynamics explain how HMGB1 lowers the persistence length of DNA and makes it more flexible (McCauley et al., 2007). It has been hypothesized that HMGB1 alters how nucleosomal DNA wraps around the octamer core (Agresti and Bianchi, 2003; Travers, 2003). The bend induced by the HMG boxes is proposed to increase unwrapping of nucleosomal DNA. Indeed, yeast (Hmo1 and Nhp6) and Drosophila (HMG-D) homologs of HMGB1 partially unwrap nucleosomal DNA (McCauley et al., 2019; Ragab and Travers, 2003). Additionally, the autoinhibition of HMGB1’s C-terminal tail is relieved by interaction with the H3 tail (Kawase et al., 2008; Ueda et al., 2004; Watson et al., 2014). It is unclear how these activities impact the competition between HMGB1 and H1 on chromatin. Further, the absence of structures of HMGB1 bound to a nucleosome has limited a mechanistic understanding of how HMGB1 increases nucleosome dynamics.

**Figure 1.**
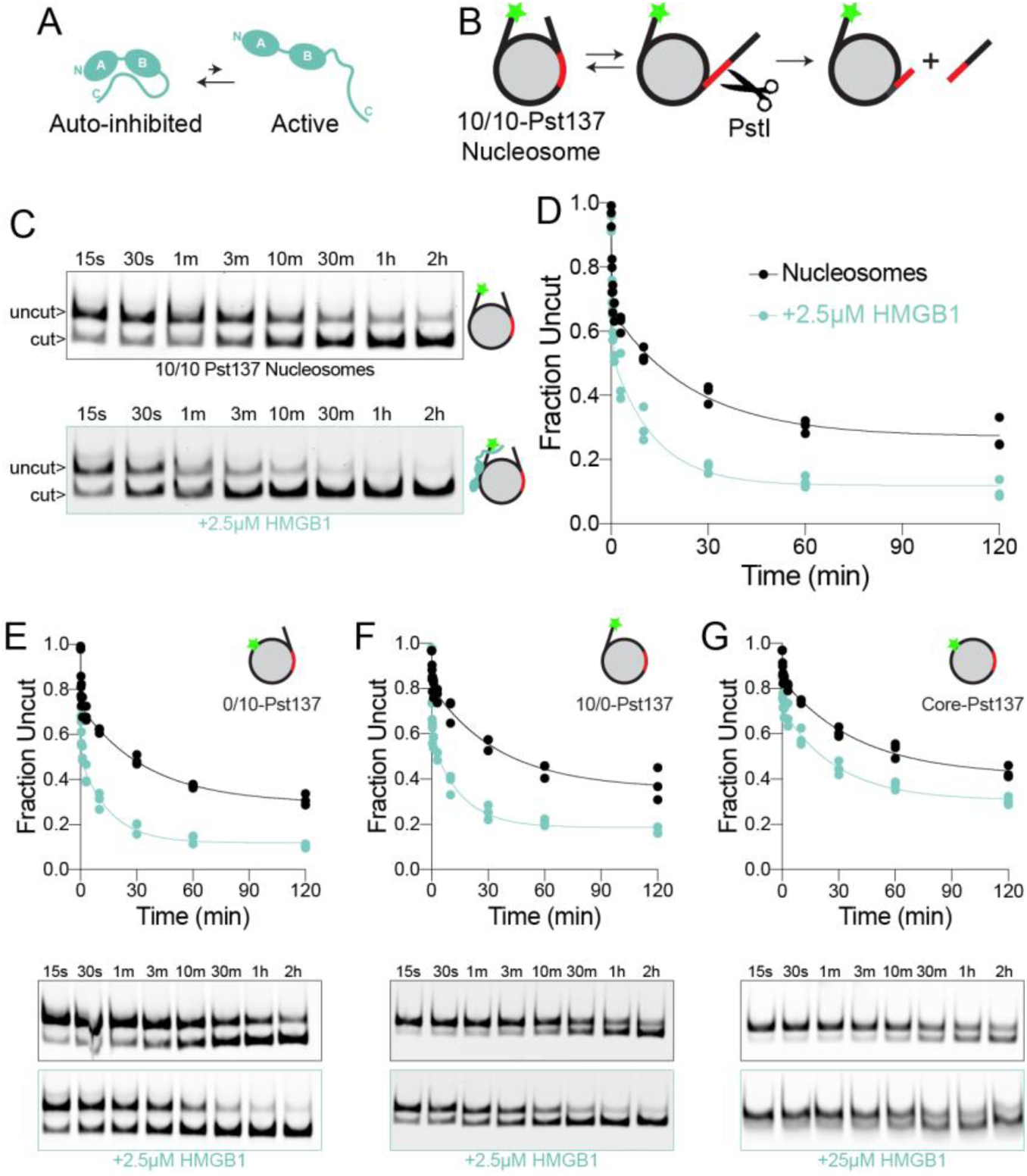
HMGB1 grants access to nucleosomal DNA. (A) A schematic of autoinhibition of HMGB1’s DNA binding A and B-boxes by the negatively charged C-terminal tail. (B) A schematic of the restriction enzyme accessibility (REA) assay showing how nucleosomal DNA cutting by the restriction enzyme PstI occurs when nucleosomal DNA transiently unwraps from the octamer core. The green star represents the fluorescent label on the end of the DNA. (C) Representative gels of time courses of REA on 10/10-Pst137 nucleosomes with (bottom) and without (top) the presence of 2.5µM HMGB1. PstI cleavage results in the appearance of the lower, cut band over time. (D) Quantification of three replicate experiments of (B). The fraction of DNA that remains uncut is plotted as a function of time. The data for all three replicates are fit by a two-step exponential decay function (E, F and G) Top panels show quantification of three replicate REA experiments on 0/10-Pst137 (E), 10/0-Pst137 (F) and Core-Pst137 (G) with (teal) and without (black) HMGB1. The fraction of DNA that remains uncut is plotted as a function of time. The data for all three replicates are fit by a two-step exponential decay function. Bottom panels show representative gels of time courses of REA on nucleosomes with (bottom) and without (top) the presence of 2.5µM HMGB1 or 25µM HMGB1 for Core Pst137.

In this study, using a quantitative assay to measure the transient unwrapping of nucleosomal DNA, and cryo-electron microscopy (cryo-EM) to determine structures of HMGB1 on nucleosomes, we find that HMGB1 and H1 do not compete for binding the same location. Rather, our data suggest that HMGB1 binds and deforms DNA at multiple sites along the nucleosomal and flanking DNA, counteracting H1’s inhibition of DNA accessibility without displacing it. We further show that HMGB1 increases the dynamics of H1 bound to condensed chromatin and of H1-bound chromatin itself. Overall, our findings provide structural clarity on how HMGB1 affects nucleosomal DNA and explain how HMGB1 and H1 counter each other’s activities to regulate chromatin dynamics at the atomic and mesoscales.

## Results

### HMGB1 grants access to nucleosomal DNA by enhancing DNA unwrapping

Previous studies have shown that HMG box-containing proteins enhance unwrapping of nucleosomal DNA (McCauley et al., 2019; Ragab and Travers, 2003). So we first sought to quantitatively test how human HMGB1 affects this process. Restriction enzyme accessibility (REA) assays have been used to show that nucleosomal DNA dynamically and partially unwraps from the octamer core (Polach and Widom, 1995). By placing a restriction enzyme site within the nucleosomal DNA, the transient unwrapping of DNA from the octamer core can be measured by the rate of restriction enzyme cutting (Figure 1B). DNA unwrapping preferentially occurs near the entry/exit sites of nucleosomal DNA, where fewer histone-DNA interactions need to be disrupted to make the DNA accessible (Anderson and Widom, 2000). Given that HMGB1 has been hypothesized to bind near the entry exit site (An et al., 1998; Nightingale et al., 1996), we used nucleosomes with 10 base pairs (bp) of flanking DNA on either side of the Widom 601 nucleosome positioning sequence, which was modified to contain a PstI restriction enzyme site centered at position 137 (10/10-Pst137, Figure 1B). We find that restriction enzyme cleavage is enhanced in the presence of HMGB1 (Figure 1C and 1D), consistent with HMGB1’s role in increasing DNA unwrapping.

It has previously been shown that HMGB1 prefers the presence of flanking DNA to interact with nucleosome substrates (Bonaldi et al., 2002; Ueda et al., 2004). To assess whether flanking DNA is also essential for HMGB1’s effect on nucleosomal DNA unwrapping, we performed REA assays on nucleosomes with asymmetric flanking DNA (0/10 or 10/0) and found that HMGB1 enhances restriction enzyme cutting on both substrates (Figure 1E and 1F). Although nucleosomal DNA is slightly less accessible when there is no flanking DNA proximal to the PstI cut site (10/0-Pst137, Figure 1F), HMGB1 still enhances restriction enzyme cutting on these nucleosomes. This suggests that HMGB1 enhances the existing unwrapping dynamics of nucleosomal DNA and that HMGB1’s binding preference for nucleosomes with flanking DNA is independent of its effect on nucleosomal DNA accessibility once bound.

These results prompted us to look more closely at how exactly HMGB1 interacts with nucleosome substrates. In electrophoretic mobility shift-based binding experiments using fluorescently end-labelled nucleosomes, HMGB1 produces two up-shifted bands (Figure S1A). The two bands are consistent with a 2:1 stoichiometry of HMGB1 on nucleosomes. Interestingly, this two-step binding pattern is observed when HMGB1 binds symmetric 10/10 and asymmetric 0/10 nucleosomes (Figure S1A), suggesting that symmetric flanking DNA is dispensable for HMGB1’s stoichiometry. Additionally, there is minimal effect on nucleosome affinity when removing one half of the flanking DNA, while removing all flanking DNA causes a 10-fold decrease in affinity (Figure S1B and S1C). Yet HMGB1 still promotes nucleosome unwrapping of core nucleosomes (Figure 1F). Overall, these results indicate that while flanking DNA increases the affinity of HMGB1 for nucleosomes, it is not essential for HMGB1’s activity on nucleosomes, suggesting the presence of other interactions that facilitate HMGB1’s activity.

It is known that the HMGB1 C-terminal tail interacts with the histone H3 N-terminal tail (Kawase et al., 2008; Ueda et al., 2004; Watson et al., 2014). It is also known that the H3 tail interacts with nucleosomal DNA and inhibits DNA accessibility (Ghoneim et al., 2021; Polach et al., 2000). Therefore, it has been hypothesized that the interaction between the HMGB1 C-terminal tail and the H3 tail could result in sequestration of the H3 tail and relief of autoinhibition by HMGB1’s C-terminal tail, both of which would increase HMGB1’s ability to unwrap DNA (Watson et al., 2014). Consistent with previous studies, we found that deleting the H3 tail significantly increases DNA unwrapping compared to wildtype nucleosomes (Figure S2A and S2B) (Ghoneim et al., 2021; Polach et al., 2000). We expected that deleting the autoinhibitory C-terminal tail of HMGB1 (HMGB1-ΔC) would also increase the DNA unwrapping activity of HMGB1. However, we found that HMGB1-ΔC has a different effect. HMGB1-ΔC does not yield the two-step nucleosome binding pattern of wildtype HMGB1 and instead binds non-specifically at a much higher stoichiometry and affinity (Figure S1A). Additionally, HMGB1-ΔC inhibits restriction enzyme cutting (Figure S2A and S2B), likely due to steric inhibition by the HMG boxes bound non-specifically to nucleosomal DNA. This suggests that the interaction between the HMGB1 C-terminal tail and the histone H3 N-terminal tail plays an additional activating role beyond relieving autoinhibition. We speculate that this additional role is to positionally localize HMGB1 near the entry/exit site, which effectively couples sequestration of the H3 tail away from nucleosomal DNA, relief of auto-inhibition by the HMGB1 C-terminal tail and local bending and unwrapping of nucleosomal DNA by the HMG boxes.

### HMGB1 transiently interacts at multiple sites along the nucleosomal DNA

To directly visualize where HMGB1 interacts with the nucleosome and how this affects nucleosomal DNA unwrapping, we collected a cryo-electron microscopy (cryo-EM) dataset of 0/10 nucleosomes bound by HMGB1 in the absence of cross-linking. Using this dataset, we determined a 3D reconstruction where we detect one HMG box bound at superhelical location (SHL) −2 on the nucleosomal DNA (Figure 2A, 2B, S3, and S4). This position is notably similar to recent cryo-EM structures of the related transcription factors SOX2 and SOX11 (Dodonova et al., 2020), and could suggest a preferred location on the nucleosomal DNA for HMG box domains. While the local resolution of the HMG box density was insufficient for de novo model building, we find that the density most likely represents box A of HMGB1 by docking previously determined models of HMGB1 box A and box B (PDB 4QR9 and PDB 2GZK, respectively) (Sánchez-Giraldo et al., 2015; Stott et al., 2006) into the cryo-EM density followed by real space refinement (Figure 1B and S6). We base this interpretation on the ability to see density for phenylalanine in box A but not in box B (Figure S6) and the fact that box A bends DNA less than box B (Teo et al., 1995), which would make it more likely to fit the curvature of nucleosomal DNA. As in the structure of SOX11, we also see a widening of the minor groove of the nucleosomal DNA at SHL −2 (Figure 2C), consistent with the effect of HMG box domains on DNA. Notably, we could only resolve a single HMG box in our structure, while there could be up to four HMG boxes if two HMGB1 molecules are bound to a single nucleosome. We attribute this observation to HMGB1’s high off rate and hypothesize that there are other regions of nucleosomal DNA where the HMG boxes can transiently interact that are being averaged out in our structure. Thus, there may be an ensemble of rapidly interconverting nucleosome bound conformations of HMGB1.

**Figure 2.**
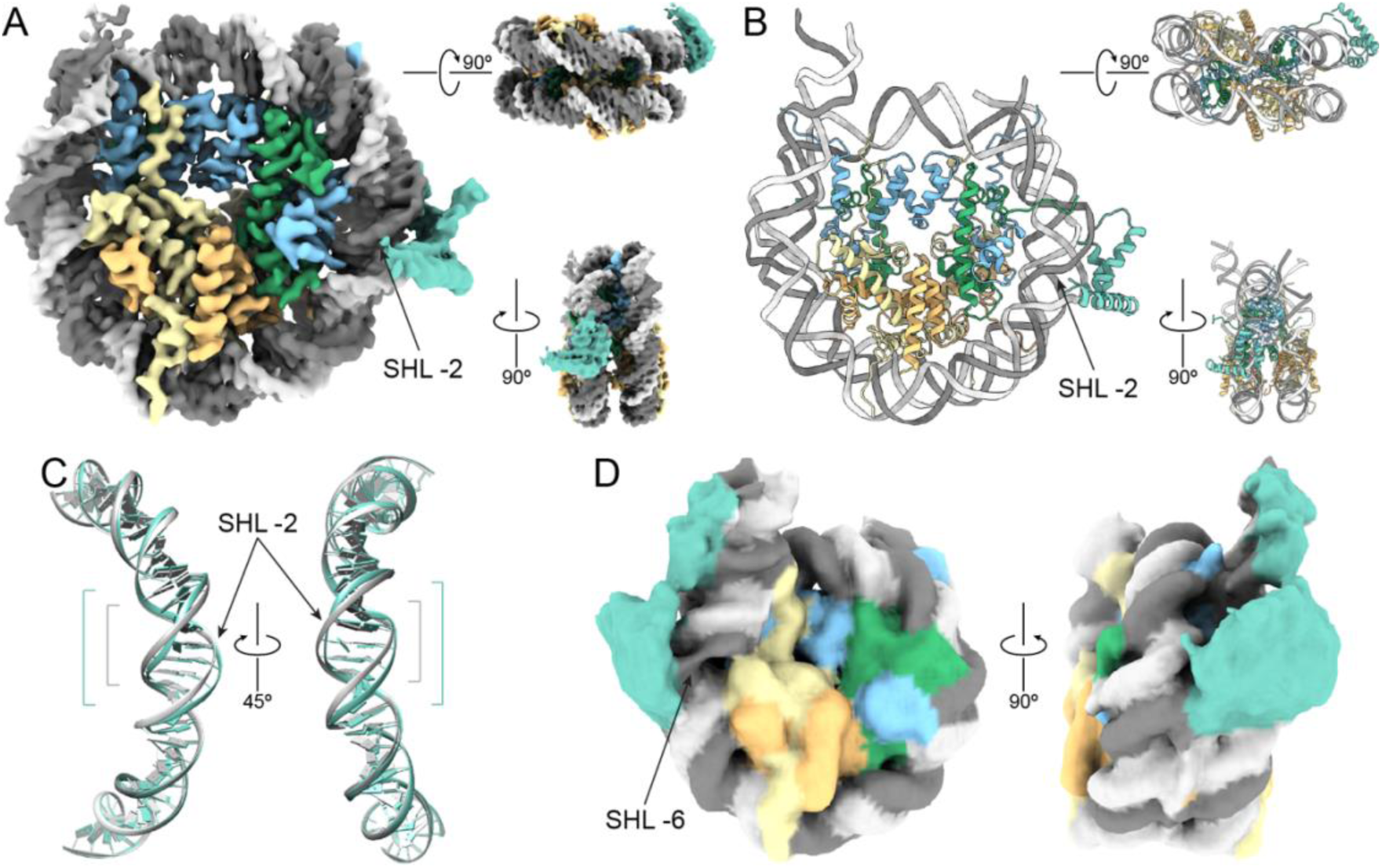
HMGB1 distorts nucleosomal DNA at multiple sites. (A) Cryo-EM density map of the A box of HMGB1 bound to a 0/10 nucleosome at SHL −2 at 2.9Å resolution. (B) Atomic model built using the density in (A). (C) A comparison of the DNA bound by HMBG1 at SHL −2 (gray) to the unbound DNA at SHL +2 (teal), showing the widening of the minor grove by the HMG box. (D) An additional cryo-EM class revealed by cryoDRGN showing extra density for HMGB1 bound at SHL −6.

To visualize other conformations of HMGB1 on nucleosomes that may be present in our dataset, we used cryoDRGN (Zhong et al., 2021). This analysis revealed various classes from our dataset, including ones that correspond to HMGB1 bound at SHL −2 as detected by our earlier analysis (Figure S5). Interestingly, in one particular class, there is extra density at SHL −6, which we interpret to be HMGB1 (Figure 2D). In this class, we see a modest kink in the nucleosomal DNA, representing the bend induced by the HMG boxes (Figure 2D and S7). Additionally, in a related reconstruction, we also see this nucleosomal DNA kink at SHL −6 even though HMGB1 density is more poorly resolved (Volume 19, Figure S5 and S7). The DNA distortions at SHL −6 in these two classes are consistent with our REA results, where we see an increase in DNA accessibility at position 137, which resides between SHL −6 and −7. Together, these structures imply that the two HMG boxes of HMGB1 bind transiently at multiple sites along the nucleosomal DNA. This finding is inconsistent with binding competition between HMGB1 and H1 as none of the binding sites we observe for HMGB1 overlap with H1’s binding site at the dyad. When we do observe binding of an HMG box, it correlates with modest local distortion of the nucleosomal DNA. Rapid binding and unbinding of the two HMG boxes and corresponding transient DNA distortion at multiple sites could contribute to HMGB1’s overall effect of increasing nucleosomal DNA accessibility. Further if the C-terminus is anchored by the H3 tail, such positioning could synergize with preferential DNA unwrapping at the entry/exit sites, where it is less restricted by contacts with the octamer core.

### HMGB1 selectively stabilizes a nucleosome conformation with exposed nucleosomal DNA

In all REA experiments, we observe that nucleosomal DNA cutting contains two kinetic phases, suggesting that nucleosomes exist in two conformations that interconvert more slowly than cutting by PstI. In this model, one of the conformations is cut more rapidly (fast-cut) than the other (slow-cut), with respective rate constants of k_fast_ and k_slow_. Interestingly, k_fast_ is similar to the rate constant of cutting naked DNA (Figure S2D and S2E), suggesting that the fast-cut conformation has nucleosomal DNA at position 137 that is fully accessible to PstI. We don’t attribute the fast cut population to contaminating free DNA as we do not detect substantial free DNA in our assembled nucleosomes (Figure S2C).

To investigate how HMGB1 influences these populations of nucleosomes, we performed REA at a range of HMGB1 concentrations and compared the rate constants of the fast and slow-cut populations (k_fast_ and k_slow_) as a function of HMGB1 concentration (Figure 3A and 3B). To our surprise, we found that the magnitude of k_fast_ or k_slow_ was not substantially dependent on the concentration of HMGB1 (Figure 3C). Instead, the fraction of nucleosomes that were cut fast increased with increasing concentrations of HMGB1, indicating that at higher HMGB1 concentrations a greater population of nucleosomes have more accessible DNA. The increase in the fast cut fraction as a function of HMGB1 concentration resembles a binding curve (Figure 3D), implying that HMGB1 binding correlates with an increase in the population of fast-cutting nucleosomes. To quantify this effect, we fit a model to the data whereby HMGB1 binds the fast-cut nucleosomes more strongly than the slow-cut nucleosomes, resulting in shift in the equilibrium towards the fast cut population when nucleosomes are fully bound by HMGB1 (Figure 3E). This analysis revealed that HMGB1 has a roughly 4-fold binding preference for fast-cut over slow-cut nucleosomes (Figure 3E). Additionally, the analysis showed that 50% of HMGB1’s stimulatory effect on nucleosomal DNA unwrapping occurred at ∼1.4 µM, a K_1/2_ value that is comparable to the K_D_ measured for HMGB1 to nucleosomes (Figure S1C). Overall, these results are consistent with a model where HMGB1 increases nucleosomal DNA accessibility by preferentially stabilizing fast-cut nucleosomes.

**Figure 3.**
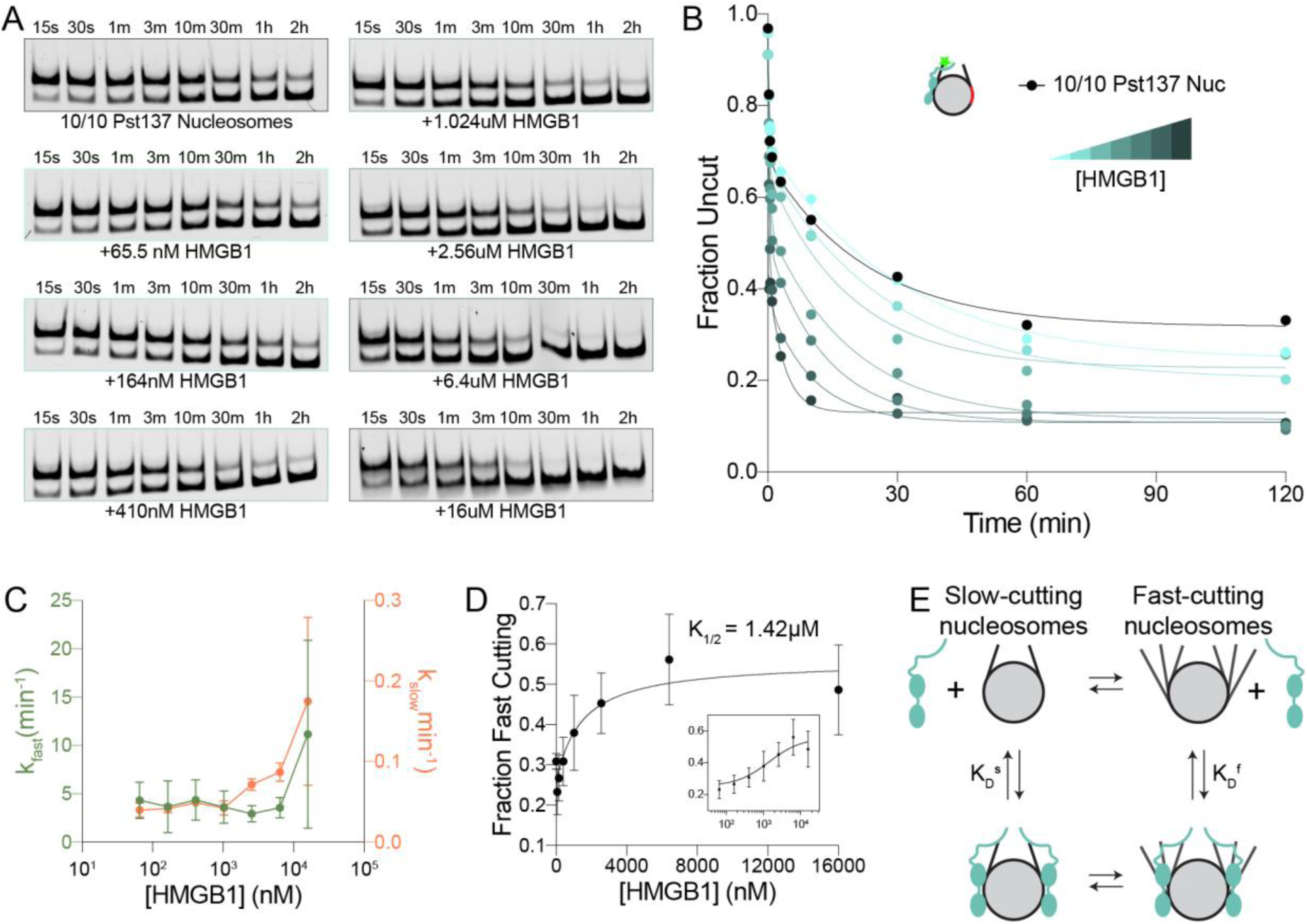
HMGB1 selectively stabilizes a conformation of nucleosomes with fast-cutting nucleosomal DNA. (A) Representative time-course gels of REA experiments on 10/10-Pst137 nucleosomes as a function of varying HMGB1 concentrations (65.5nM, 164nM, 410nM, 1.024µM, 2.56µM, 6.4µM, and 16µM). (B) Quantification of REA experiments in (A). Each time course is fit by a two-step exponential decay. (C) Rate constants (k_fast_ and k_slow_) for experiments in (A) are shown as a function of HMGB1 concentration. Error bars represent the standard deviation for three separate replicates. (D) The fraction of nucleosomes that are described by k_fast_ is plotted as a function of HMGB1 concentration. Error bars represent the standard deviation for three separate replicates. Curve represents a fit by an equation derived from the model in (E). An overall half-maximal concentration of HMGB1 for its effect is 1.42µM (K_1/2_). Inset shows the same plot with a logarithmic x-axis. (E) A schematic of a model showing a slow and a fast-cutting population of nucleosomes that interconvert slowly on timescale of PstI cutting and are bound by HMGB1 with different affinities. K_D_ ^s^ is 2.4µM while K_D_ ^f^ is 650nM. K_D_’s are derived from the equations in Materials and Methods.

### HMGB1 and H1 obstruct each other’s effects on nucleosomes

To directly test how HMGB1 and H1 compete to regulate nucleosomal DNA accessibility, we performed REA experiments in the presence of both proteins on a minimal nucleosome substrate for H1 binding (10/10-Pst137) (White et al., 2016). Consistent with previous REA results using *Drosophila* H1 (Ragab and Travers, 2003), when nucleosomes are bound by H1 there is no detectable cutting (Figure 4A), as a closed conformation of the entry/exit DNA is stabilized by H1 (Syed et al., 2010). However, even when HMGB1 is added at concentrations 16-fold over its K_D_ to nucleosomes bound by stoichiometric H1, cutting is not recovered (Figure 4A and 4B). H1’s K_D_ for nucleosomes is in the picomolar-nanomolar range, compared to the micromolar range K_D_ for HMGB1, and this large difference in affinities could in principle explain why HMGB1 cannot outcompete H1 (White et al., 2016). However, this explanation does not take into account that the two proteins can interact with each other via their oppositely charged C-terminal tails (Cato et al., 2008). We therefore hypothesized that this direct interaction enables H1’s inhibition of HMGB1 activity. To test this possibility, we investigated the effects of a mutant of H1 (H1-ΔCTE), which lacks the C-terminal basic tail that interacts with HMGB1. This construct still inhibits DNA unwrapping in our REA assay (Figure 4A). However, the inhibition of unwrapping is rescued by HMGB1 in a concentration-dependent manner (Figure 4A and 4B). We interpret this result to mean that direct interaction between the two proteins via their C-terminal tails is essential for H1’s inhibition of HMGB1. An alternative explanation is that deletion of H1’s C-terminal tail reduces its affinity for nucleosomes ∼100-fold (White et al., 2016), such that HMGB1 is now able to compete for binding at lower concentrations.

**Figure 4.**
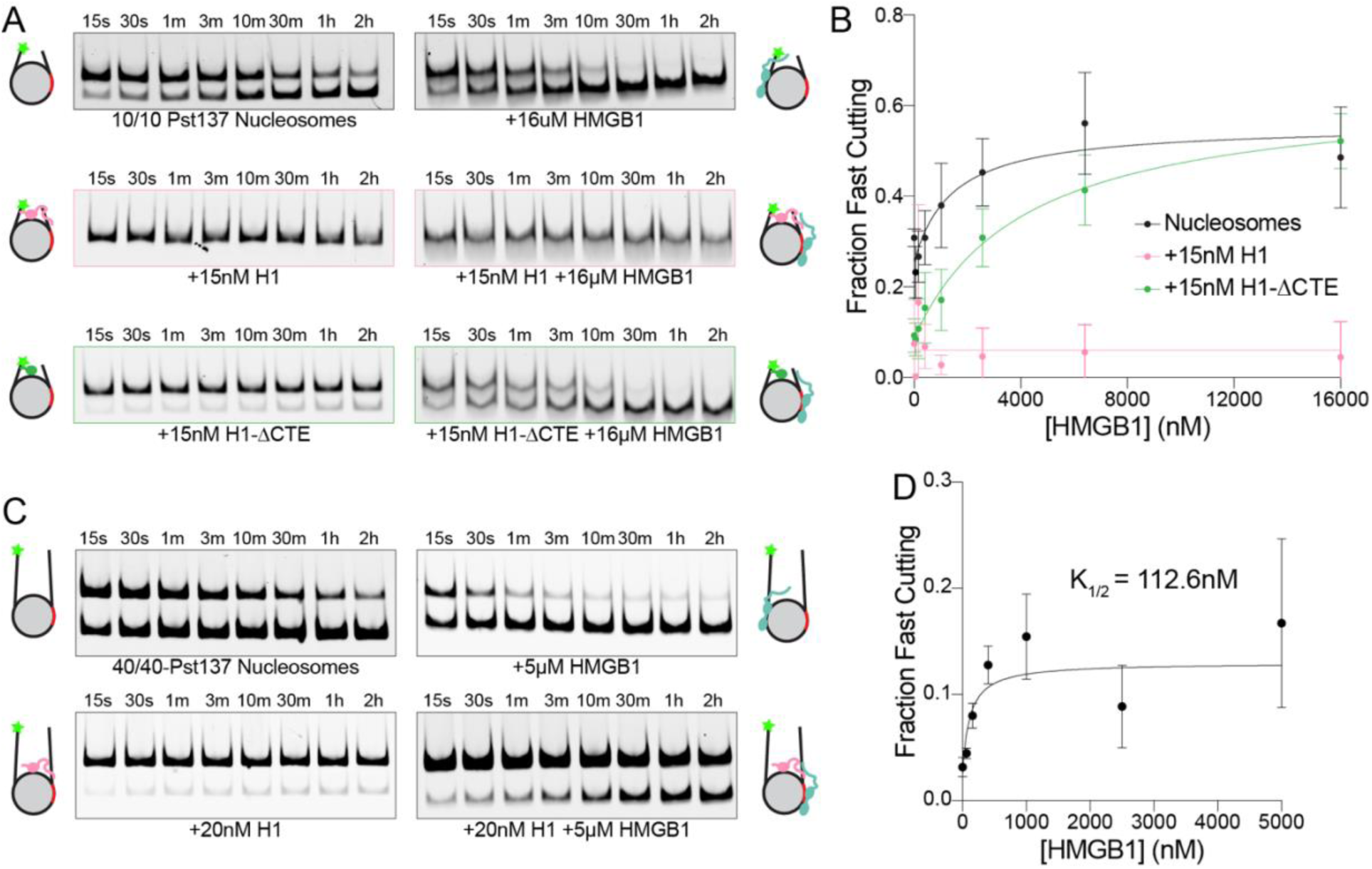
HMGB1 and H1 compete to tune nucleosomal DNA accessibility. (A) Representative time-course gels of REA experiments in the presence of indicated concentrations of H1 and/or HMGB1. (B) The fraction of fast cutting nucleosomes is shown as a function of HMGB1 concentration. Before HMGB1 is added, nucleosomes are either bound by no H1 (black), 15nM H1 (coral), or 15nM H1-ΔCTE (green). Curve represents a fit by an equation derived from the model in Figure 3E. (C) Representative REA time courses of 40/40-Pst137 nucleosomes with or without H1 and/or HMGB1. (D) The fraction of fast cutting nucleosomes is shown as a function of HMGB1 concentration. 40/40-Pst137 nucleosomes were pre-incubated with 20nM H1 before HMGB1 was added. Curve represents a fit by an equation derived from the model in Figure 3E. K_1/2_ is 112.6nM, K_D_ ^s^ is 126.3nM and K_d_ ^f^ is 21.25nM.

In mammalian chromatin, linker lengths vary from 5bp to over 100 bp (Abdulhay et al., 2020). To investigate the HMGB1 and H1 interplay in the context of linker lengths longer than 10 bp, we used nucleosomes with 40bp of flanking DNA on either side (40/40-Pst137). Interestingly, basal restriction enzyme cutting is enhanced in the presence of longer flanking DNA (Figure S2F), suggesting the length of flanking DNA influences the accessibility of nucleosomal DNA. Similar to the 10/10 nucleosomes, H1 also inhibits REA on 40/40 nucleosomes (Figure 4C). However unlike 10/10 nucleosomes, on 40/40 nucleosomes, HMGB1 can partially rescue the inhibition of DNA unwrapping caused by H1 indicating that linker DNA length also tunes the competition between

HMGB1 and H1 (Figure 4C). Even in the presence of saturating HMGB1, the rescue of nucleosomal DNA accessibility is partial (Figure 4C and 4D). This result indicates that HMGB1 counteracts the effects of H1 without displacing H1 from nucleosomes, implying that HMGB1 and H1 can co-occupy a nucleosome. Additionally, the concentration at which DNA unwrapping is 50% rescued (K_1/2_) is 112.6nM (Figure 4D), similar to HMGB1’s K_D_ for DNA and 40/40 nucleosomes (Figure S1C). This finding suggests that HMGB1 binding to 40/40 nucleosomes is insensitive to the presence of H1, further implying the formation of a ternary complex. Unlike on 10/10 nucleosomes, the flanking DNA on 40/40 nucleosomes is sufficient to accommodate an entire HMGB1 molecule. We therefore propose that 40/40 nucleosome allow more modes of HMGB1 binding thereby changing how HMGB1 and H1 interact and compete.

### HMGB1 stimulates nucleosomal DNA accessibility within chromatin

We next asked whether HMGB1 promotes access to nucleosomal DNA in the context of a chromatin template. To look at accessibility of DNA within chromatin, we employed the single-molecule adenine methylated oligonucleosome sequencing assay to test chromatin accessibility on assembled templates (SAMOSA-ChAAT) (Abdulhay et al., 2023). We assembled nucleosomes onto a DNA template containing 12 successive 601 sequences interspersed by 46bp of flanking DNA (Gibson et al., 2019). Looking at the pattern of DNA accessibility within the chromatin array, we noticed increased and asymmetric accessibility near the entry/exit sites of the nucleosome (Figure 5A), consistent with transient nucleosomal DNA unwrapping and the asymmetry of the 601 sequence (Anderson and Widom, 2000; Ngo et al., 2015). In response to increasing amounts of HMGB1, DNA within the chromatin arrays becomes more exposed (Figure 5A). Furthermore, we see that the increase in DNA accessibility is most pronounced near the entry/exit sites of the nucleosome resulting in a decreased nucleosome footprint size (Figure 5B and 5C), consistent with the model that HMGB1 enhances existing unwrapping dynamics to expose nucleosomal DNA. The increase in restriction enzyme cutting we observe at position 137 with nucleosomes, is well within the region of increased accessibility we observe by SAMOSA-ChAAT on chromatin arrays, suggesting that HMGB1 acts similarly on mononucleosomes and nucleosomes within chromatin arrays.

**Figure 5.**
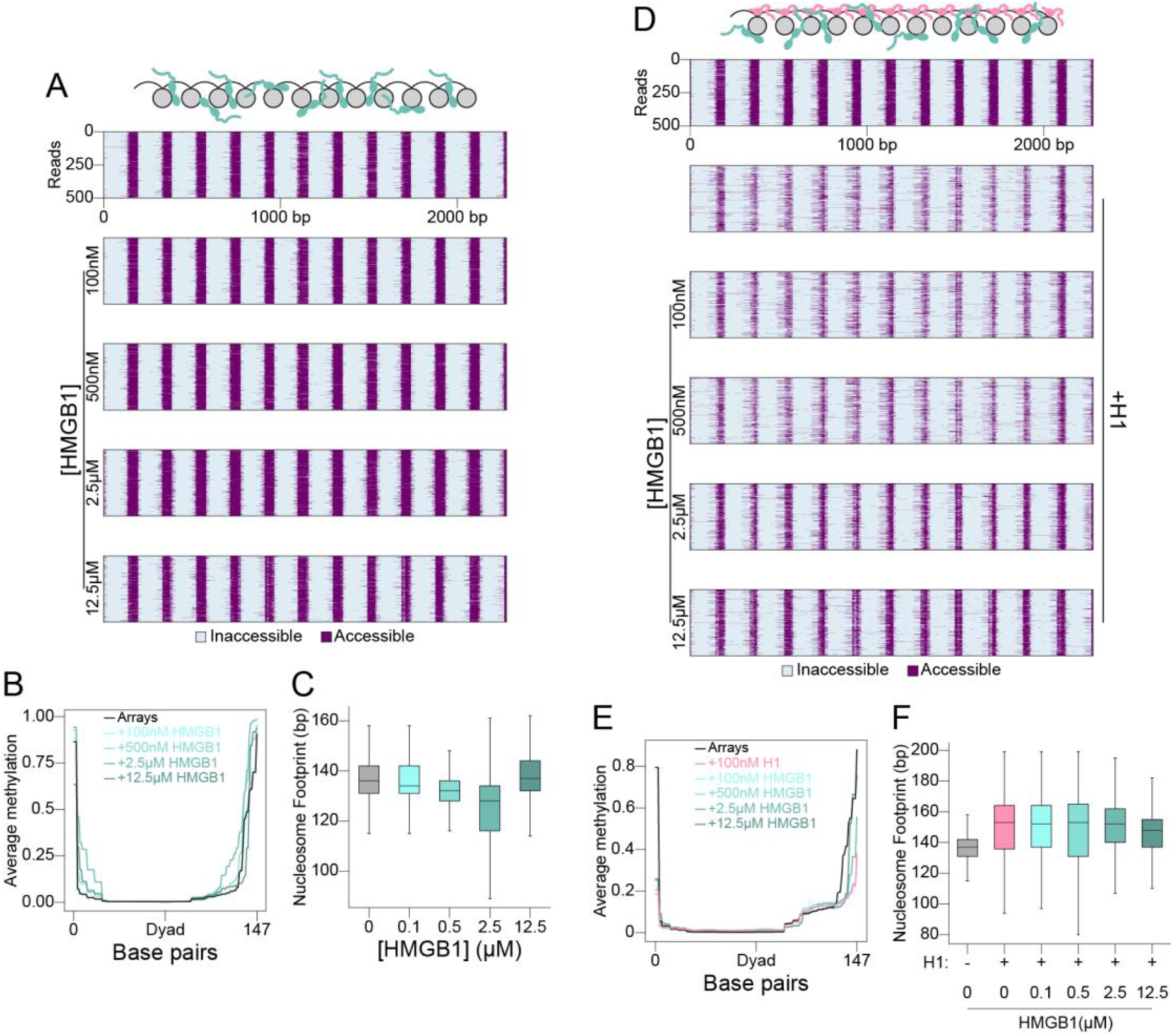
HMGB1 increases accessibility of DNA within chromatin. (A) Heat maps of SAMOSA-ChAAT experiments showing nucleosome footprints on a 12x601 array with a range of HMGB1 concentrations. Purple represents DNA that is accessible as assessed by the presence of adenine methylation. Grey represents DNA that is inaccessible as assessed by the lack of methylation and is interpreted as a nucleosome footprint. (B) The average methylation across every 601 sequence within the array is combined to show an average nucleosome footprint within the array. (C) Box and whiskers plot of nucleosome footprint size is plotted as a function of HMGB1 concentration. Solid line represents the median footprint size, while the height of the box represents the interquartile range, and the error bars represent the farthest data point within 1.5x the interquartile range from the box. (D) Heat maps of SAMOSA-ChAAT experiments showing nucleosome footprints on a 12x601 array with H1 and a range of HMGB1 concentrations. Purple and grey represent accessible and inaccessible DNA, respectively as in (A). (E) The average methylation across every 601 sequence within the array is combined to show an average nucleosome footprint within the array. (F) Box and whiskers plot of nucleosome footprint size is plotted as a function of HMGB1 concentration. Solid line represents the median footprint size, while the height of the box represents the interquartile range, and the error bars represent the farthest data point within 1.5x the interquartile range from the box.

Interestingly, at the highest concentration of HMGB1 tested, we notice a decrease in accessibility and a larger footprint (12.5µM HMGB1, Figure 5A, 5B and 5C). We attribute this effect to bending of the linker DNA caused by binding of additional HMGB1 molecules leading to internucleosome interactions that inhibit access to nucleosomal DNA. This possibility is consistent with previous observations showing that yeast Hmo1 and Nhp6 can promote compaction of chromatin arrays (McCauley et al., 2019).

To explore how HMGB1 and H1 compete to regulate DNA accessibility within chromatin, we performed SAMOSA-ChAAT on chromatin arrays in the presence of both proteins. Consistent with H1’s role in forming stable, higher-order structures of chromatin arrays (Robinson et al., 2006; Song et al., 2014), when we add H1 to our arrays, the amount of DNA protection increases (Figure 5D). Consistent with REA results on 40/40 nucleosomes, when increasing amounts of HMGB1 are added in the presence of H1, we see a corresponding rescue of DNA accessibility near the entry/exit sites (Figure 5D and 5E). This is also evidenced by a decrease in the average nucleosome footprint size at higher HMGB1 concentrations (Figure 5F). This result indicates that the opposing effects of HMGB1 and H1 observed on mononucleosomes translate to a chromatin substrate.

### HMGB1 makes H1 and condensed chromatin more dynamic

Chromatin arrays also provide the opportunity to test the dynamics of HMGB1 and H1 in the context of condensed chromatin, which may more accurately recapitulate *in vivo* chromatin properties. Cellular data on the competition between HMGB1 and H1 comes from fluorescence recovery after photobleaching (FRAP) studies, which showed that fluorescently labelled H1 recovers faster in the presence of increased concentrations of HMGB1 (Catez et al., 2004). However, it is unclear if this effect is due to direct binding competition between HMGB1 and H1 or an indirect effect. To investigate direct effects of HMGB1 on H1’s binding and turnover dynamics on condensed chromatin *in vitro*, we generated chromatin condensates. We compacted the chromatin arrays by the addition of 3mM MgCl_2_ and 100mM KCl in the presence or absence of Alexa555-labeled H1, such that H1 to nucleosome stoichiometry is 1:1, and varying concentrations of HMGB1. HMGB1 causes chromatin condensates containing H1 to become larger, although we see no loss of H1 intensity within condensates (Figure 6A and 6B). These data suggest that HMGB1 does not simply displace H1 by occupying H1’s binding site and are consistent with our data suggesting co-occupancy of HMGB1 and H1 on nucleosomes. We next explored whether HMGB1 affects the dynamics of H1 within condensed chromatin.

**Figure 6.**
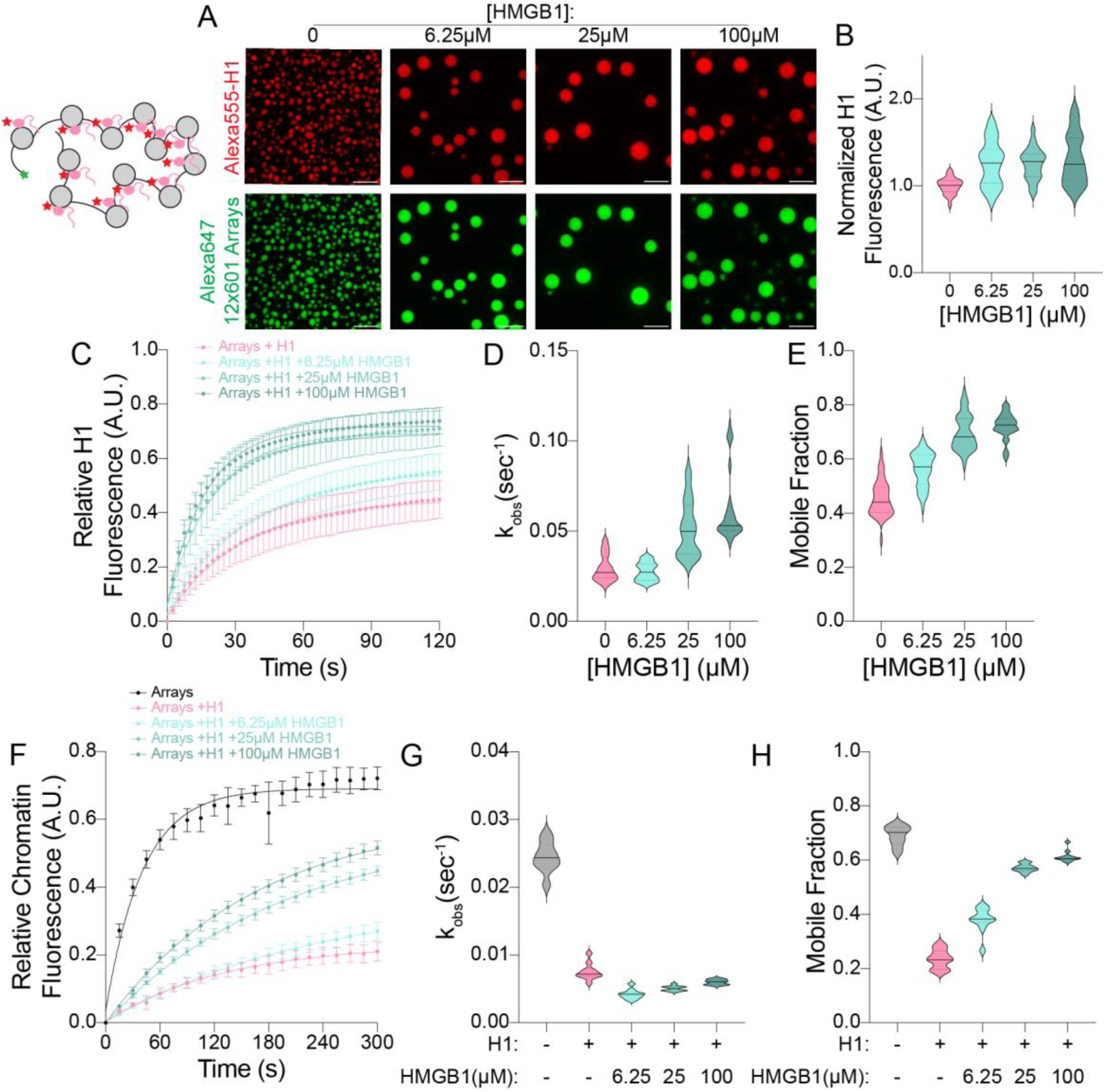
HMGB1 increases the turnover dynamics of H1 and chromatin. (A) Representative microscopy images of chromatin condensates. 30nM Alexa647-12x601 arrays are mixed with 360nM Alexa555-H1 and varying concentrations of HMGB1. Scale bar represents 10µm. (B) Violin plots of the quantification of the average Alexa555-H1 fluorescence intensity within chromatin condensates. All values are normalized to chromatin and H1 alone with no HMGB1 added. Solid and dotted lines represent the median and interquartile values respectively. Values represent an average of three experimental replicates with n>10 condensates per replicate. (C) Quantification of the fluorescence recovery after photobleaching of Alexa555-H1 within chromatin condensates. Values are normalized from 0 to 1 corresponding to post and pre-bleach respectively. Points represent an average of three experimental replicates with n>10 condensates per replicate. Error bars represent standard deviation of all condensates within the three replicates. Data are fit to a one-phase exponentially association (D and E) Violin plots of the quantification of the fits to the FRAP data in (C). The rate constant for recovery (k_obs_) and the plateau of recovery (Mobile Fraction) are plotted for the four different conditions in (D) and (E) respectively. Solid and dotted lines represent the median and interquartile values respectively. (F) Quantification of the fluorescence recovery after photobleaching of Alexa647-12x601 arrays within condensates. Values are normalized from 0 to 1 corresponding to post and pre-bleach respectively. Points represent a single experimental replicate with n>10 condensates. Error bars represent standard deviation of all condensates. Data are fit to a one-phase exponentially association. (G and H) Violin plots of the quantification of the fits to the FRAP data in (F). The rate constant for recovery (k_obs_) and the plateau of recovery (Mobile Fraction) are plotted for the five different conditions in (G) and (H) respectively. Solid and dotted lines represent the median and interquartile values respectively.

Consistent with previous results (Gibson et al., 2019), we observe that H1 fluorescence recovers to about 45% of pre-bleach intensity within two minutes (Figure 6C). Upon addition of HMGB1, we observe an increase in the rate of recovery and the mobile fraction of H1 that recovers, both of which are HMGB1 concentration-dependent (Figure 6D and 6E). Taken together with the observation that HMGB1 does not decrease the amount of H1 in condensates, we conclude that HMGB1 alters the on/off dynamics of H1’s interaction with chromatin, but that binding of HMGB1 and H1 to chromatin is not mutually exclusive.

This model raises the possibility that while HMGB1 and H1 do not compete to bind the same site within chromatin, their activities on regulating chromatin dynamics are in competition with one another. To investigate this possibility, we performed FRAP on the Alexa647-labeled chromatin within condensates. Consistent with previous FRAP results within chromatin condensates (Gibson et al., 2019), we find that 70% of the chromatin recovers to pre-bleach intensity within five minutes (Figure 6F). The addition of H1 slows recovery by ∼3-fold and reduces the fraction that recovers to 25%, also consistent with previous results (Gibson et al., 2019) (Figure 6F, 6G, and 6H). Upon addition of increasing amounts of HMGB1, we see a partial rescue of chromatin dynamics (Figure 6F). HMGB1 restores the mobile fraction of chromatin to 60% but interestingly does not increase the rate constant of recovery (Figure 6G and 6H). This result suggests that HMGB1’s effect is due to altering H1’s activity since in the absence of H1, HMGB1 has minimal effect on chromatin dynamics (Figure S8). Together with results showing that H1 is not displaced from the droplets, these findings provide further evidence that HMGB1 and H1 modulate each other’s activities by co-occupying chromatin.

## Discussion

Compared to its competitor H1, less is known about HMGB1’s interaction with nucleosomes and its effects on chromatin at the mesoscale. Our findings suggest that HMGB1 enhances the dynamics of nucleosomes through rapid sampling and local distortion of multiple locations on nucleosomal DNA by its HMG boxes while its C-terminus is anchored by the H3 tail. Our results also shed light on the competition between HMGB1 and H1. Whereas H1 inhibits dynamics of nucleosomes and chromatin, HMGB1 appears to act as a molecular stir bar to increase chromatin dynamics at multiple scales. This interplay between two abundant nuclear architectural proteins presents new opportunities for chromatin regulation. Below we discuss the mechanistic and biological implications of our findings in the context of previous results.

### Multiple modes of engagement explain how HMGB1 enhances nucleosome dynamics

Substantial previous work has led to the model that HMGB1 exposes nucleosomal DNA through its DNA bending activity (Agresti and Bianchi, 2003; Balliano et al., 2017; Bonaldi et al., 2002; Hepp et al., 2014; Joshi et al., 2012; Travers, 2003; Ugrinova et al., 2009). Work with the H3 tail peptide indicates that interactions between the C-terminal tail of HMGB1 and the H3 tail relieve auto-inhibition to allow bending of nucleosomal DNA and facilitate nucleosomal DNA accessibility (Watson et al., 2014). Here we’ve directly tested this model in the context of a nucleosome. Our results show that the HMG boxes directly interact with and deform nucleosomal DNA. Additionally, our results uncover an additional role for the interaction between HMGB1’s C-terminal tail and the H3 tail. Not only does this interaction sequester the H3 tail away from nucleosomal DNA, allowing DNA to unwrap more readily and relieve autoinhibition of the HMG boxes as proposed previously (Watson et al., 2014), but in addition our results imply that the H3 tail orients HMGB1 on nucleosomes in a way that promotes unwrapping of nucleosomal DNA (Figure 7A). We propose that interaction of the C-terminal tail of HMGB1 with the H3 tail spatially localizes the HMG boxes to certain regions of nucleosomal DNA, such as SHL −2 and SHL −6. Amongst these sites, those where nucleosomal DNA is less restrained by histone interactions, HMG box induced DNA bending is more likely to result in unwrapping. Further, we speculate that HMGB1’s high off rate and the dynamic interconversion between bound and unbound states allow the nucleosome to rapidly sample multiple states in which different regions of nucleosomal DNA become accessible. Substantial previous work has uncovered how histone chaperones catalyze interconversions between assembled and disassembled nucleosomes and how ATP-dependent chromatin remodelers slide and conformationally alter nucleosomes (Clapier et al., 2017; Winkler and Luger, 2011). The mechanism for HMGB1 provides a third and qualitatively distinct way to make nucleosomal DNA accessible.

**Figure 7.**
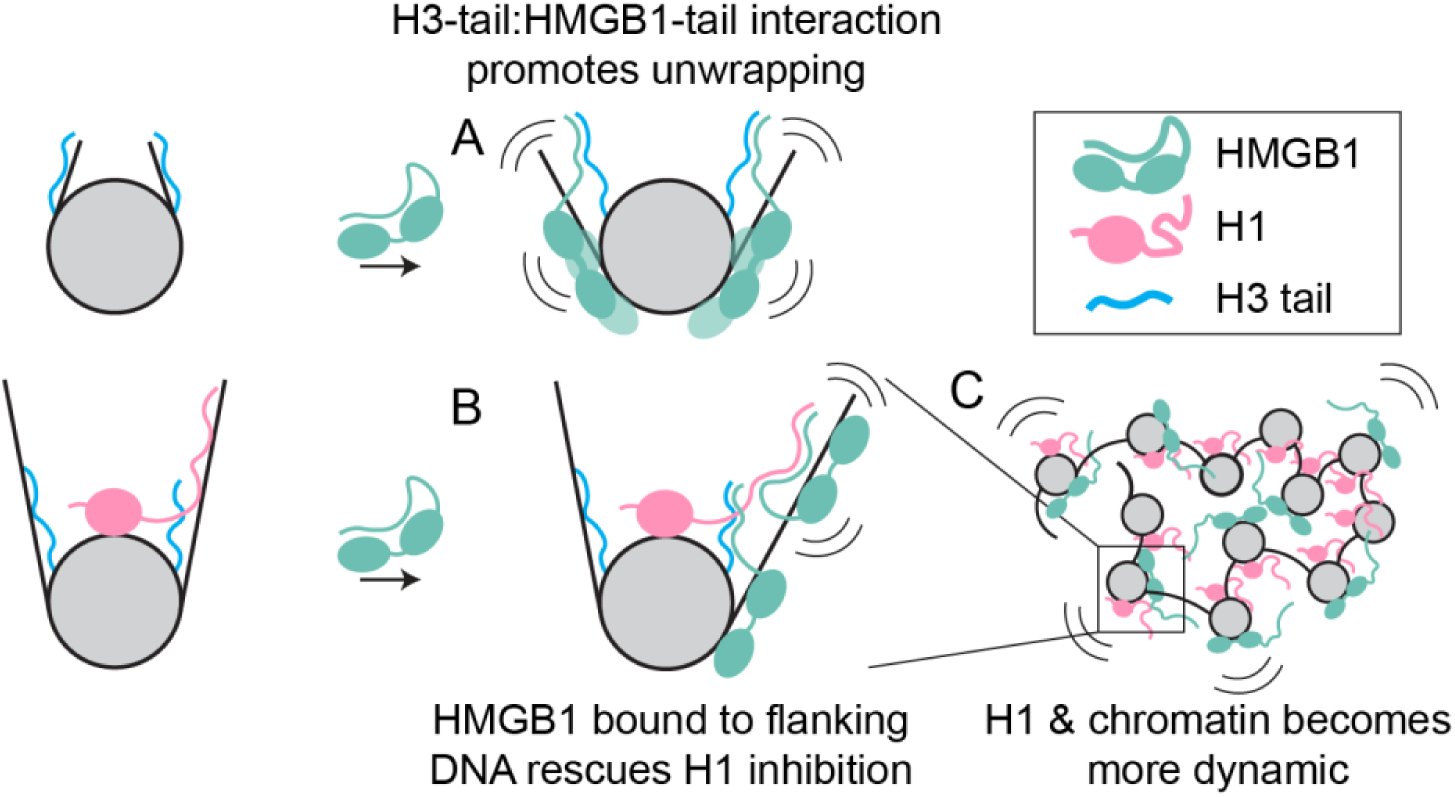
HMGB1 and H1 compete to regulate chromatin dynamics across scales. (A) HMGB1 promotes unwrapping through interaction with the H3 tail. This replaces the H3 tail’s interaction with nucleosomal DNA and relieves autoinhibition of HMGB1, allowing the HMG boxes to bind and unwrap nucleosomal DNA. (B) When longer flanking DNA is present, HMGB1 is able to rescue H1 inhibition of DNA unwrapping. The extra flanking DNA allows additional HMGB1 molecules to interact, which loosens H1’s interaction with the flanking DNA, allowing HMGB1 to rescue DNA unwrapping. (C) HMGB1’s loosening of H1’s interaction with flanking DNA makes both H1 and chromatin itself more dynamic within condensed chromatin.

### HMGB1 and H1 oppose one another’s activities as a function of flanking DNA length

Our data suggest that HMGB1 and H1 counteract each other’s activities while co-occupying a nucleosome. Further, the ability of HMGB1 to counteract H1’s inhibition of DNA unwrapping increases with increasing flanking DNA length and when the H1 C-terminal tail is deleted. H1 is proposed to reduce nucleosome dynamics by interacting with and stabilizing the flanking DNA proximal to the entry/exit site (Syed et al., 2010). It is also proposed that interaction between the HMGB1 C-terminal tail and the H1 C-terminal tail relieves autoinhibition of HMGB1 (Cato et al., 2008). We hypothesize that on nucleosomes with short flanking DNA (10 bp or less) HMGB1 can interact with the H1 C-terminal tail, but that this interaction does not sufficiently outcompete the interaction of the H1 tail with nucleosomal DNA. As a result, on nucleosomes with 10 bp of flanking DNA, HMGB1 cannot overcome H1’s inhibition of DNA unwrapping. However, when longer flanking DNA is present, the H1 C-terminal tail can now bind flanking DNA in addition to nucleosomal DNA and HMGB1 molecules can be accommodated on the flanking DNA in addition to the core nucleosome (Figure 7B). We propose that in this context the C-terminal tails of HMGB1 and H1 interact resulting in (i) stabilization of HMG box binding to flanking DNA, (ii) displacement of the H1 tail from flanking DNA and (iii) bending of the flanking DNA. Additionally, there are likely cooperative effects from the distortion of the flanking DNA that could translate to HMGB1 molecules bound to nucleosomal DNA. Together these coupled outcomes would increase the probability of DNA unwrapping. In this model, the tail-tail interaction between HMGB1 and H1 and the length of flanking DNA tune the two proteins’ effects on DNA accessibility. Overall, we propose that the transient interactions of HMGB1, and the H1 and H3 tails with nucleosomal and flanking DNA creates an ensemble of rapidly interconverting nucleosome bound states. This allows for rapid switching between different states in response to changes in linker length and HMGB1 and H1 concentrations to regulate DNA accessibility.

Deletion of H1’s C-terminal tail increases H1’s turnover *in vivo* (Phair et al., 2004). Therefore, our model predicts that HMGB1 would increase H1’s turnover dynamics within condensed chromatin by loosening the H1 C-terminal tail’s interaction with flanking DNA, which is what we observe. This interaction would also have the effect of rescuing H1’s inhibition of chromatin dynamics, since the H1 tail is necessary for its inhibition of chromatin turnover (Gibson et al., 2019). HMGB1 thus has the effect of a molecular stir bar, enhancing the dynamics of chromatin bound proteins and chromatin at the level of nucleosomes and condensed chromatin.

### Mechanistic Basis for HMGB1’s diverse roles in vivo

Overexpression of HMGB1 has been linked to transcription activation (Aizawa et al., 1994), suggesting HMGB1 reduces the inhibitory effect of nucleosomes. Interestingly, a knockout of HMGB1 in mice is associated with a global reduction in nucleosomes, and *in vitro*, HMGB1 enhances the assembly of chromatin (Celona et al., 2011; Lange et al., 2008; Osmanov et al., 2013). Our findings reconcile these seemingly contradictory results. We observe that HMGB1 drives a transient increase in nucleosomal DNA accessibility and distorts nucleosomal DNA without disassembling the nucleosome. We therefore propose that HMGB1-induced nucleosomal DNA flexibility allows nucleosomes to dynamically accommodate transient structural changes in DNA, without falling apart. Such accommodation would increase nucleosome stability in the face of factors that try to access nucleosomal DNA, while also increasing DNA access. Indeed, the FACT nucleosome chaperone complex, that protects nucleosomes from polymerases, contains an HMG box domain (Winkler and Luger, 2011).

Consistent with the proposed model, at the nucleosome scale HMGB1 has been shown to enhance the affinity of the estrogen receptor to a nucleosomal site and enhance ATP-dependent remodeling by ISWI and SWI/SNF family of chromatin remodelers (Bonaldi et al., 2002; Hepp et al., 2014; Joshi et al., 2012; Patenge et al., 2004; Ugrinova et al., 2009). HMGB1 has also been shown to increasing binding of various transcription factors (TF’s) to DNA via DNA bending and direct interactions with the TF (Boonyaratanakornkit et al., 1998; Das et al., 2004; Decoville et al., 2000; Jayaraman et al., 1998; Joshi et al., 2011; Oñate et al., 1994; Zappavigna et al., 1996; Zwilling et al., 1995). We therefore speculate that a transient HMGB1-TF complex could act as bi-partite pioneer factor enabling access to nucleosomal DNA sequence without disassembly of the histone octamer. Indeed, the SOX family of pioneer factors contains HMG boxes that have evolved sequence specificity, hinting at a possible universal mechanism of HMG box domains for accessing nucleosomal DNA.

Our findings further suggest that the H3 tail-HMGB1 interaction not only releases auto-inhibition by the HMGB1 C-terminal tail, but also orients the HMG boxes for effective DNA unwrapping. Post-translation modifications of the H3 tail could thus alter how HMGB1 interacts with nucleosomes and its effect on DNA accessibility and serve as cues to where in the genome HMGB1 acts. One possibility is that HMGB1 cooperates with histone acetyltransferases to induce DNA accessibility. Consistent with this model, HMGB1 localizes to sites in the genome marked by acetylation of Lys27 on H3 (Sofiadis et al., 2021).

### Consequences of competition between HMGB1 and H1 *in vivo*

In vivo, where measured, HMGB1 is found at lower concentrations than H1 (Catez et al., 2004; Duguet and de Recondo, 1978). Yet, the dynamics of H1 *in vivo* are sensitive to HMGB1 concentrations indicating that HMGB1 can counteract the effects of H1 despite binding nucleosomes at least 1000-fold more weakly than H1 (Bonaldi et al., 2002; Ueda et al., 2004; White et al., 2016). Our model provides one explanation for how HMGB1 effectively counteracts H1 activity. In this model, HMGB1 concentrations do not need to be high enough to competitively displace H1 from a nucleosome. HMGB1 concentrations only have to be high enough to co-occupy nucleosomes in a manner that allows the C-terminal tail of HMGB1 to interact with the H1 tail. In early studies of rat liver cells, HMGB1 was estimated at 10^6^ molecules per nucleus, compared to approximately 10^7^ H1 molecules and 10^7^ nucleosomes (Catez et al., 2004; Duguet and de Recondo, 1978). Given average nuclear volumes (Maul and Deaven, 1977), HMGB1 exists in the nucleus at 5-10µM, which is close to the half maximal concentrations that we measure for HMGB1’s effect on tuning H1-bound chromatin dynamics. In stem cells where HMGB1 expression is highest, H1 turnover is also increased (Meshorer et al., 2006; Müller et al., 2004), suggesting that HMGB1’s stir bar activity may be especially important for pluripotency. HMGB1 expression is also elevated in some cancers, which could present an opportunity for therapeutic intervention (Müller et al., 2004).

Along with expression levels, phosphorylation of the H1 C-terminal tail could then modulate the competition between HMGB1 and H1. Indeed in metaphase, when H1 phosphorylation is higher, HMGB1 is displaced from chromosomes while H1 remains bound (Contreras et al., 2003; Falciola et al., 1997). H1 phosphorylation has also been linked to both activation and repression of transcription (Bhattacharjee et al., 2001; Dou et al., 1999; Dou and Gorovsky, 2000), suggesting that there could be a variety of modes of regulation of this competition via H1 phosphorylation. Further exploration of the role of H1 tail phosphorylation will reveal how competition with HMGB1 is modulated *in vivo*.

Recent attempts to determine nucleosome structures *in situ* have found that the majority of nucleosomes do not have stable flanking DNA conformations, and many nucleosomes appear partially unwrapped (Arimura et al., 2021; Cai et al., 2018c, 2018b, 2018a; Tan et al., 2023). We propose that the competition between HMGB1 and H1 is responsible for creating some of these more heterogeneous nucleosome structures, yet HMGB1, due to its high off rate and multiple modes of binding cannot be captured structurally, unlike H1. Additional heterogeneity may also arise from variations in linker lengths as we find that flanking DNA length is an important factor that tunes the competition between HMGB1 and H1, with H1 effects predominating at shorter linker lengths and HMGB1 effects at longer lengths. Linker DNA lengths *in vivo* vary from 5bp to over 100bp (Abdulhay et al., 2020) and thus likely regulate where in the genome HMGB1 or H1 affect nucleosomal DNA accessibility.

This study emphasizes the essential role of architectural proteins in modulating global effects on chromatin dynamics. Despite what the name suggests, we find that the abundant architectural protein HMBG1 does not serve as a structural support for chromatin, but rather our data suggests that HMBG1 increases the dynamics and accessibility of chromatin to enable DNA access without nucleosome disassembly. Further we propose that the interplay between HMGB1 and architectural proteins like H1 that act more canonically to inhibit chromatin dynamics increases the number of states of the system allowing chromatin dynamics to be finely tuned at the molecular and mesoscale.

## Acknowledgements

We thank Julia Tretyakova for purification of all histones used in this manuscript. We thank Michael K. Rosen for the H1.4 and H1.4-ΔCTE expression plasmids. We thank SoYeon Kim for tireless help and advice on condensate microscopy experiments. We thank the UCSF Center for Advanced Light Microscopy and the Nikon Imaging Center for access to tools and expertise for microscopy experiments. We thank John D Gross and Danica Galonić Fujimori for thorough advice throughout this work and helpful comments on preparation of the manuscript. We thank all members of the Narlikar Lab for stimulating discussion of this project at all stages. This work was supported by grants from the National Institutes of Health to G.J.N. (R35GM127020), to V.R. (DP2-HG012442), to G.J.N. and V.R. (U01-DK127421), to Y.C. (R35GM140847), and to U.S.C. (F32GM137463). Equipment at the UCSF cryo-EM facility was partially supported by the NIH (grants S10OD020054, S10OD021741, and S10OD025881). Y.C. is an investigator at the Howard Hughes Medical Institute. Portions of this work were funded through the gracious support of the UCSF Program for Breakthrough Biomedical Research and the Sandler Fellows program.

## Author Contributions

Conceptualization, H.S.S. and G.J.N.; Methodology, H.S.S. and G.J.N.; Software, C.M.M. and V.R.; Formal Analysis, H.S.S., U.S.C., and C.M.M.; Investigation, H.S.S., U.S.C., and C.M.M.; Writing – Original Draft, H.S.S. and G.J.N.; Writing – Review & Editing, H.S.S., U.S.C., C.M.M., V.R., Y.C., and G.J.N.; Supervision, G.J.N.

## Supplemental Information

Supplemental Figures S1-S10 and Table S1

## Methods

### Preparation of DNA substrates

DNA used for mononucleosomes was generated using large scale PCR of a plasmid containing the Widom 601 sequence. Various primers, labeled with either Alexa Fluor 488 or Cy5 (IDT), were used to generate different lengths of flanking DNA around the 601 sequence. When noted in the text, the 601 sequence contained a restriction enzyme site for PstI at position 137. PCR products were run over an 8% acrylamide gel and the band was cut out based on the DNA shadow. The gel band was crushed by passage through a 5-mL syringe and soaked in 1x TE overnight before the crushed gel was removed with a 0.22-µm filter. The DNA was further ethanol precipitated and resuspended in 1x TE.

DNA for chromatin arrays was generated from a plasmid containing 12 successive 601 sequences, interspersed by 46 bp’s of linker DNA (Gibson et al., 2019). The plasmid was grown in STBL2 cells (Thermo Fisher) and giga prepped (QIAGEN). The plasmid was then digested with EcoRV to cut out the array sequence and digest the plasmid backbone. To make Alexa Fluor 647-labeled arrays for condensate experiments, the DNA was additionally digested with XhoI to leave a sticky end on one end of the DNA. The array DNA was then purified over a Sephacryl S-1000 column and fractions containing the array DNA were pooled and isopropanol precipitated, before being resuspended in 1x TE. We note that two successive runs over the S-1000 column were necessary to fully remove the plasmid backbone.

To generate labeled array DNA, the sticky end generated by XhoI digest was filled in using Klenow polymerase (Invitrogen) with an Alexa Fluor 647-labeled dCTP (Invitrogen) along with unlabeled dATP, dTTP, and dGTP according to manufacturer’s instructions. DNA was then purified by phenol:chloroform extraction and ethanol precipitation before resuspension in 1x TE.

### Assembly of nucleosomes and chromatin arrays

Recombinant *Xenopus laevis* histones H3, H4, H2A, and H2B were expressed and purified from *E. coli* as previously described (Luger et al., 1999). Histone octamer and dimer was then reconstituted as previously described (Luger et al., 1999). Mononucleosome and chromatin arrays were assembled via salt gradient dialysis as previously described (Luger et al., 1999). Mononucleosomes were further purified over a glycerol gradient. All samples were quantified using a NanoDrop to measure the concentration of DNA within the sample.

### HMGB1 purification

A codon-optimized cDNA of *Homo sapiens* HMGB1 was generated by Twist Biosciences and cloned into a pBH4 plasmid backbone. Briefly the construct contains an N-terminal 6xHis tag followed by a TEV protease cleavage site, which when cleaved leaves a single Gly-Ser linker before the start Met of HMGB1. The plasmid is transformed into Rosetta (DE3) competent cells (Sigma Aldrich) and grown in 2xYT (20g tryptone, 10g NaCl, 10g yeast extract per 1L) supplemented with 50µg/mL carbenicillin and 25µg/mL chloramphenicol at 37°C. Once cells have reached log phase (OD = 0.5-0.6), expression was induced with 1mM IPTG at 37°C for 4 hours. Cells were then pelleted by spinning at 5000 x g for 30min at 4°C and pellets were stored at −80°C.

Pellets were resuspended in Lysis Buffer with 2x protease inhibitor (20mM HEPES-KOH pH 7.5, 500mM KCl, 10% Glycerol, 7.5mM Imidazole, 2mM BME, 2mM PMSF, 2µg/mL Pepstatin A, 6µg/mL Leupeptin, and 4µg/mL Aprotinin). The cell suspension was homogenized by passage through a Dounce homogenizer. Cells were lysed with an Emulsiflex at 15,000 psi for 15 minutes. Lysate was then clarified by centrifugation at 40,000 x g for 30min at 4°C. Lysate was then incubated with TALON affinity resin (Takara, 1mL bead volume/1L bacterial culture) for 1hour at 4°C while nutating. Beads were then washed in batch two times with two bead volumes of Lysis Buffer with 1x protease inhibitor (20mM HEPES-KOH pH 7.5, 500mM KCl, 10% Glycerol, 7.5mM Imidazole, 2mM BME, 1mM PMSF, 1µg/mL Pepstatin A, 3µg/mL Leupeptin, and 2µg/mL Aprotinin) to remove any remaining lysate. Beads were then added to plastic disposable gravity columns and washed with 20 bead volumes of Lysis Buffer (20mM HEPES-KOH pH 7.5, 500mM KCl, 10% Glycerol, 7.5mM Imidazole, 2mM BME). His-tagged HMGB1 was then eluted with 5 bead volumes of Elution Buffer (20mM HEPES-KOH pH 7.5, 150mM KCl, 400mM Imidazole, 2mM BME) (Figure S10A). Cleavage of the His tag was then induced with TEV protease (150µg TEV/1L bacterial culture) and left to dialyze overnight in TEV dialysis buffer (20mM HEPES-KOH, 150mM KCl, 2mM DTT) (Figure S10B).

TEV-cleaved, full-length HMGB1 was then added to a Mono-Q anion exchange column equilibrated with 10% Buffer QB (Buffer QA: 20mM HEPES-KOH pH 7.5, 3mM DTT Buffer QB: 20mM HEPES-KOH pH 7.5, 1.5M KCl, 3mM DTT). HMGB1 was eluted from the Mono-Q column with a linear gradient from 10 to 60% Buffer QB (150mM to 900mM KCl) over 20 column volumes. Full length HMGB1 comes off between 32-35% Buffer QB (∼500mM KCl), while many products that do not contain the full C-terminal domain come off at lower [KCl], therefore this anion exchange step is crucial to obtaining pure, full-length HMGB1 (Figure S10C and S10D).

These fractions are then pooled and concentrated in a 3000 Da molecular weight cut off spin filter device. HMGB1 is finally purified over a Superdex 75 Increase size exclusion column equilibrated in Size Exclusion buffer (20mM HEPES-KOH pH 7.5, 300mM KCl, 10% Glycerol, 3mM DTT) (Figure S10E and S10F). Fractions containing HMGB1 are pooled and concentrated in a 3000 Da molecular weight cut off spin filter device, before being aliquoted, flash-frozen, and stored at −80°C in Size Exclusion buffer.

HMGB1-ΔC is purified identically, except a HiTrap SP cation exchange column is used in place of the Mono-Q anion exchange column, as the pI of HMGB1-ΔC is basic, while HMGB1-WT is acidic. A 10 to 60% gradient of Buffer QB (150mM to 900mM KCl) is still used, and HMGB1-ΔC elutes between 29-33% Buffer QB (∼465mM KCl). We do not see any contaminating expression products, because the C-terminal domain has been removed.

### H1 purification

*Homo sapiens* H1.4 purification was adapted from (Gibson et al., 2019) using a construct containing an N-terminal MBP, followed by a TEV cleavage site, followed by H1.4 (full length or ΔCTE), followed by an additional TEV cleavage site, and a C-terminal 6xHis tag. This construct was transformed into Rosetta pLysS competent cells (EMD Millipore) and grown in 2xYT (20g tryptone, 10g NaCl, 10g yeast extract per 1L) supplemented with 50µg/mL carbenicillin and 25µg/mL chloramphenicol at 37°C. Once cells have just reached log phase (OD = 0.4), cultures are moved to 18°C for 1 hour. After 1 hour, expression is induced with 0.5mM IPTG for 18 hours at 18°C. Cultures are then pelleted at 5000 x g for 30min at 4°C and pellets are stored at −80°C.

Pellets are resuspended in Lysis Buffer with 2x protease inhibitor (20mM HEPES-KOH pH 7.5, 1M NaCl, 10% Glycerol, 7.5mM Imidazole, 2mM BME, 2mM PMSF, 2µg/mL Pepstatin A, 6µg/mL Leupeptin, and 4µg/mL Aprotinin) The cell suspension was homogenized by passage through a Dounce homogenizer. Cells were lysed with an Emulsiflex at 15,000 psi for 15 minutes. Lysate was then clarified by centrifugation at 40,000 x g for 30min at 4°C. Lysate was then incubated with TALON affinity resin (Takara, 1.5mL bead volume/1L bacterial culture) for 1hour at 4°C while nutating. Beads were then washed in batch two times with two bead volumes of Lysis Buffer with 1x protease inhibitor (20mM HEPES-KOH pH 7.5, 1M NaCl, 10% Glycerol, 7.5mM Imidazole, 2mM BME, 1mM PMSF, 1µg/mL Pepstatin A, 3µg/mL Leupeptin, and 2µg/mL Aprotinin) to remove any remaining lysate. Beads were then added to plastic disposable gravity columns and washed with 20 bead volumes of Lysis Buffer with 1x protease inhibitor (20mM HEPES-KOH pH 7.5, 1M NaCl, 10% Glycerol, 7.5mM Imidazole, 2mM BME, 1mM PMSF, 1µg/mL Pepstatin A, 3µg/mL Leupeptin, and 2µg/mL Aprotinin). Tagged H1 was then eluted with 5 bead volumes of Elution Buffer (20mM HEPES-KOH pH 7.5, 150mM NaCl, 10% Glycerol, 350mM Imidazole, 2mM BME, 1mM PMSF, 1µg/mL Pepstatin A, 3µg/mL Leupeptin, and 2µg/mL Aprotinin). Eluted protein was then incubated with amylose affinity resin (NEB, 2mL bead volume/1L bacterial culture) for 1hour at 4°C while nutating. The beads are then added to plastic disposable gravity columns and washed with 20 bead volumes of Amylose Wash Buffer (20mM HEPES-KOH pH 7.5, 150mM NaCl, 10% Glycerol, 2mM BME, 1mM PMSF, 1µg/mL Pepstatin A, 3µg/mL Leupeptin, and 2µg/mL Aprotinin). H1 is then eluted with 5 bead volumes of Amylose Elution Buffer (20mM HEPES-KOH pH 7.5, 150mM NaCl, 10% Glycerol, 2mM BME, and 1% Maltose). Cleavage of the MBP and His tags was then induced with TEV protease (225µg TEV/1L bacterial culture) and left to dialyze overnight in TEV dialysis buffer (20mM HEPES-KOH, 150mM KCl, 2mM DTT).

H1 is then loaded onto a HiTrap SP cation exchange column equilibrated with TEV dialysis buffer. H1 is then eluted with a linear gradient from 10 to 60% Buffer SB (Buffer SA: 20mM HEPES-KOH pH 7.5, 2mM BME Buffer SB: 20mM HEPES-KOH pH 7.5, 1.5M NaCl, 2mM BME) over 20 column volumes. Fractions containing H1 were then incubated with 2mL bead volume of TALON affinity resin (Takara) for 1 hour at 4°C nutating, to remove contaminating His-tagged TEV protease. The beads were removed using a plastic gravity column and the flow through is concentrated using a 3000 Da molecular weight cut off spin filter device. H1 was then purified over a Superdex 75 Increase size exclusion column equilibrated in Size Exclusion buffer (20mM HEPES-KOH pH 7.5, 150mM NaCl, 10% Glycerol, 1mM DTT). Fractions containing H1 were pooled and concentrated in a 3000 Da molecular weight cut off spin filter device, before being aliquoted, flash-frozen, and stored at −80°C in Size Exclusion buffer.

For purification of fluorescently labeled H1, a Lys-Cys-Lys was added N-terminal to the start Met of H1. The protein was purified identically to H1-WT, however, before size exclusion chromatography, H1 was dialyzed overnight into Labeling Buffer (20mM HEPES-KOH pH 7.5, 150mM NaCl, 1mM TCEP). Protein was then labeled with 3-fold molar excess of Alexa Fluor 555 maleimide (Invitrogen) at room temperature for 15 minutes. The labeling was quenched with 10-fold excess DTT. The protein was then concentrated using a 3000 Da molecular weight cut off spin filter device, and the unincorporated dye was removed by purification over a Superdex 75 Increase size exclusion column equilibrated in Size Exclusion buffer (20mM HEPES-KOH pH 7.5, 150mM NaCl, 10% Glycerol, 1mM DTT). Fractions containing H1 were pooled and concentrated in a 3000 Da molecular weight cut off spin filter device, before being aliquoted, flash-frozen, and stored at −80°C in Size Exclusion buffer.

### Restriction enzyme accessibility (REA) assays

For all restriction enzyme accessibility assays, 10nM of fluorescently labeled nucleosomes containing a PstI site at position 137 within the nucleosomal DNA are used. Purified HMGB1 or H1 is added to nucleosomes in the presence of a 10x buffer that supplements to a final buffer of: 19mM HEPES-KOH pH7.5, 1mM Tris-HCl pH7.4, 30mM KCl, 45mM NaCl, 5mM MgCl_2_, 2mM DTT, 9% Glycerol, 0.11mM EDTA, 20µg/mL BSA, 0.015% Triton-X100, and 0.02% NP-40.

Nucleosomes are allowed to equilibrate with HMGB1 or H1 for 30 minutes at 37°C. When both proteins are present, H1 is allowed to equilibrate first for 30 minutes, before adding HMGB1 and waiting another 30 minutes to start the reaction. The time course is started upon addition of PstI (NEB, 100U/uL) to a final concentration of 10U/uL. At each time point, 5µL of the reaction is removed and added to a 2x REA Stop Mix (20mM Tris-HCl pH8, 70mM EDTA, 2% SDS, and 20% Glycerol). After the time course, 2uL of Proteinase K (NEB, 800U/mL) is added to each time point and allowed to incubate for 30 minutes at 55°C. For reactions in which >5µM of HMGB1 was added, an additional 2uL of 800U/mL Proteinase K was added after 15 minutes. After incubation, each time point is loaded onto a 8% acrylamide (29:1 acrylamide:bis) 1x TBE gel and run at 150V for at least 2 hours. Gels are imaged using a Typhoon laser-scanning gel imager (GE). The fraction of DNA that is uncut is measured using ImageJ and plotted as a function of time. All nucleosome REA experiments are fit to a two-step exponential decay function in GraphPad Prism:

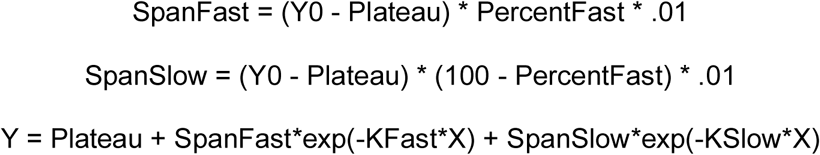

The fraction of nucleosomes that is cut fast is calculated using the following equation, which also factors in the fraction of nucleosomes that are not cut (Plateau):

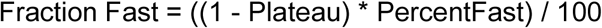

All REA experiments on DNA are fit to a one-step exponential decay function in GraphPad Prism:

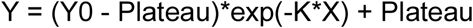

### Two-phase REA fitting and modeling

A model for how HMGB1 interacts with both fast and slow-cutting nucleosome populations is shown in Figure 3E. The fraction fast calculated above is plotted as a function of HMGB1 concentration and fit to the following equations derived based on this thermodynamic cycle to calculate the K_D_ for the fraction that is cut fast and slow (K_D_ ^f^ and K_D_ ^s^ respectively) and the overall half maximal concentration of HMGB1 (K_1/2_):

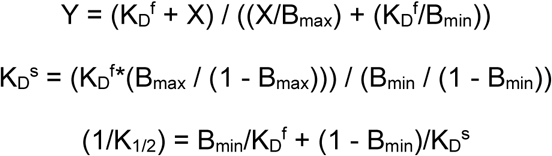

B_min_ and B_max_ represent the fraction of fast cutting nucleosomes when fully unbound (B_min_) and fully bound (B_max_).

### Electrophoretic mobility shift assays (EMSA)

Fluorescently labeled nucleosomes (10nM) were incubated with various concentrations of HMGB1 in the presence of a 10x buffer that supplements the final buffer composition to be: 20mM HEPES-KOH pH7.5, 75mM KCl, 10% Glycerol, 1mM EDTA, 2mM DTT, and 0.02% NP-40. Binding was allowed to equilibrate at room temperature for 30 minutes before loading onto a 6% acrylamide (29:1 acrylamide:bis) 0.5x TBE gel. The gel was run for 2.5 hours at 125V and imaged using a Typhoon laser scanning gel imager (GE). The fraction of nucleosomes that are bound is calculated by measuring the unbound nucleosomes band and subtracting that value from the intensity of the whole lane. The fraction bound is then plotted as a function of HMGB1 concentration and fit to the following binding equation with the Hill coefficient N:

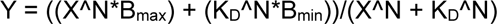

### Cryo-EM sample preparation and data collection

HMGB1 and 0/10 nucleosomes were dialyzed into EM buffer (20 mM HEPES-KOH pH 7.5, 25 mM KCl, 1 mM EDTA, 2 mM DTT, 1.5% glycerol) overnight. Due to the relatively low affinity (micromolar range), HMGB1 was mixed with nucleosomes at final concentrations of 12 µM HMGB1 and 1 µM nucleosome and allowed to incubate at 20°C for 30 minutes prior to plunge freezing on Quantifoil R 1.2/1.3 200 mesh Au grids. Grids were cleaned beforehand for 15s using a PELCO easiGLOW glow discharge system.

Sample grids were plunge frozen using a FEI Mark IV Vitrobot set at 4°C and 100% humidity. 3 µL of sample was applied to each grid. Each grid was blotted with humidity-saturated Whatman 1 filter papers for 3s with a blot force of −1 before being plunge frozen into liquid ethane.

Sample grids were first screened using a 200 kV FEI Talos Arctica at UCSF. A larger dataset was then collected on two grids on a 300 kV Titan Krios at UCSF using a K3 camera at a nominal magnification of 105kx (0.834 Å/pix). All data was collected using Serial EM v3.7 or newer.

### Cryo-EM data processing

Raw movies were motion-corrected using UCSF MotionCor2 v1.4.1 (Zheng et al., 2017). Dose-weighted micrographs were imported into cryoSPARC v4.1.1 (Punjani et al., 2017) and patch CTF estimation (multi) was used to estimate defocus values. Micrographs were filtered using Curate Exposures based on average defocus, estimated CTF, and relative ice thickness. A nucleosome map was used to generate templates for particle picking using Template Picker. A total of ∼8.5 million particles were extracted at 288 pix box downsampled to 72 pix. To remove junk particles, extracted particles were filtered through three rounds of Heterogeneous Refinement, where in each round a good nucleosome map and 3 junk maps (generated using Ab Initio on a small subset of remaining particles) were used as templates. After classification, the remaining good particles were re-extracted at 288 pix box downsampled to 192 pix and duplicate picks were removed using Remove Duplicates, resulting in 561,888 particles.

The 561,888 particles were then used in Ab Initio to generate 3 classes, and the output volumes were used as templates for Heterogeneous Refinement to further classify the remaining particles. One good class containing 339,268 particles was further refined first using Homogenous Refinement and then Non-Uniform Refinement, optimizing for per-particle defocus and per-group CTF parameters. The resulting map showed a hint of density for an HMG box at SHL −2.

To further classify the particles for nucleosomes that contain an HMG box, the particles were exported using UCSF pyem v0.5 (Asarnow et al., 2019) to RELION v4.0 (Scheres, 2012). 3D classification without alignment was performed using a spherical mask centered on the SHL −2 region. One class out of 4 showed density for the HMG box domain, and the 47,813 particles in this class were imported back into cryoSPARC v4.1.1 and re-extracted at 288 pix box. One round of 2D classification was performed to remove remaining obvious junk, resulting in 42,115 particles. This final set of particles was refined using Non-Uniform Refinement, resulting in the final map at ∼3 Å global resolution for an HMGB1 box domain bound to nucleosome at SHL −2.

### cryoDRGN analysis

The 339,268 particles that showed a hint of density for an HMG box at SHL −2 were re-extracted at 300 pix downsampled to 240 pix. The particles were exported using UCSF pyem v0.5 to generate a .star file, and a corresponding .mrcs particle stack was generated using RELION. Inputs were preprocessed using cryoDRGN v1.1.2 (Zhong et al., 2021) to generate pose and CTF .pkl files compatible with cryoDRGN. Images were downsampled further from 240 pix to 120 pix and used to train a cryoDRGN model with an 8-dimensional latent variable over 50 epochs of training. cryoDRGN analysis was then performed using the resulting model to visualize the latent space and generate density maps using default settings.

### Model building

The ∼3 Å global resolution output volume from cryoSPARC Non-Uniform Refinement was used to build a model for the HMGB1 box domain bound to nucleosome at SHL −2. The nucleosome from PDB 8V4Y (Chio et al., 2024) was used as the initial template for the nucleosome, and an HMGB1 Box A domain from PDB 4QR9 (Sánchez-Giraldo et al., 2015) was used as the initial template for the HMG box domain. DNA, histones, and the HMG box were adjusted using COOT v0.9.6 to account for differences in base pairs and residues that are resolved or not resolved in the current map. The ISOLDE plug-in (Croll, 2018) for UCSF ChimeraX v1.7.1 was used to correct for Ramachandran and rotamer outliers. COOT v0.9.6 was then used to correct bond angle and bond length outliers. The quality of all refined models was assessed using model validation in Phenix v1.18.2 and the wwPDB validation server.

### Single-molecule adenine methylated oligonucleosome sequencing assay to test chromatin accessibility on assembled templates (SAMOSA-ChAAT)

Unlabeled chromatin arrays (1ng/µL) were incubated with varying concentrations of HMGB1 and H1 (50nM when present) in the presence of 1x CutSmart Buffer (NEB), 2mM DTT and 1mM S-Adenosyl methionine in 100µL. Binding was allowed to equilibrate for 30 minutes at 37°C. In the cases when HMGB1 and H1 were present, H1 was allowed to bind first for 30 minutes before addition of HMGB1 and an additional 30 minutes to equilibrate. To induce DNA methylation, 1µL of EcoGII (NEB) was added to a final concentration of 50U/mL. Methylation proceeded for 30 minutes at 37°C before adding 10µL of 10% SDS 2.5µL ProK (NEB, 800U/mL). Proteinase K digestion proceeded for 2 hours at 65°C. Methylated DNA was purified from these reactions via 1X SPRI Select Beads.

#### PacBio Library Preparation and Sequencing

Entire binding reactions were used as input for PacBio SMRTbell library preparation. SMRTbell preparation of libraries was done using the SMRTbell prep kit 3.0 and included DNA damage repair, end repair, SMRTbell ligation, and exonuclease cleanup according to the manufacturer’s instruction. After exonuclease cleanup and purification via 1x v/v SMRTbell cleanup beads, DNA concentration was measured by Qubit High Sensitivity DNA Assay (1 µl each sample). Data was collected over 30-hour Sequel II movie runs with 2 hours pre-extension time and 2.1 polymerase.

#### SMRT Data Processing

Sequencing reads were processed as homogenous samples as described in (Abdulhay et al., 2023) with slight variations.

Chromatin Sample Processing

Raw sequencing reads from chromatin samples were processed using software from Pacific Biosciences:

1. Generate circular consensus sequences (CCS)

CCS were generated for each sequencing cell using ccs 6.9.99. The --hifi-kinetics flag was used to generate kinetics information (interpulse duration, or IPD) for each base of each consensus read. Values were stored for each base as 50*(mean logIPD) + 1.
2. Demultiplex consensus reads

Consensus reads were demultiplexed using lima. The flag ‘–same’ was passed as libraries were generated with the same barcode on both ends. This produces a BAM file for the consensus reads of each sample.
3. Align consensus reads to the reference genome

pbmm2, the pacbio wrapper for minimap2 (Li, 2018), was run on each CCS BAM file (the output of step 2) to align reads to the reference sequence, producing a BAM file of aligned consensus reads.

#### Extracting interpulse duration measurements

The IPD values were accessed from the aligned, demultiplexed consensus BAM files. Model Training

Neural network, SMM, and SVD models were trained on fully methylated and unmethylated controls similarly to (Abdulhay et al., 2023) but using IPD values from the aligned, demultiplexed consensus BAM files.

The Hidden Markov model was structured similarly to (Abdulhay et al., 2023) but was refactored from pomegranate to use cython and numba.

#### Processed data analysis

All processed data analyses and associated scripts are available at GitHub. All analyses were computed using python. Plots were constructed via Matplotlib. Each analysis is briefly described below:

#### Defining inaccessible regions and counting nucleosomes

Inaccessible regions were called from HMM output data identically to (Abdulhay et al., 2020). Briefly, inaccessible regions were defined as continuous stretches with accessibility ≤0.5.

Periodic peaks were observed that approximated sizes of regions containing one, two, three, or more nucleosomes. Cutoffs for each size were manually defined using the histogram of inaccessible region lengths from a control sample.

### Fluorescence recovery after photobleaching (FRAP) of chromatin condensates

Purified HMGB1 and H1 were dialyzed overnight in Protein Dialysis Buffer (20mM HEPES-KOH pH7.5, 200mM KCl, 7mM MgCl_2_, and 2mM BME). Arrays were diluted to 60nM in buffer from the chromatin array assembly (20mM HEPES-KOH pH7.5, 1mM EDTA, and 2mM BME). For each reaction, 10µL of arrays, 2µL H1 (or buffer), and 8µL HMGB1 (or buffer) were added to a microcentrifuge tube so that the final buffer is 20mM HEPES-KOH pH7.5, 100mM KCl, 3.5mM MgCl_2_, 0.5mM EDTA, and 2mM BME. After 30 minutes at room temperature, reactions were transferred to a 384-well glass bottom plate (Cellvis), which had been mPEGylated and passivated with BSA according to (Gibson et al., 2019). Condensates were allowed to settle in the well for at least 1 hour before imaging with a 100x oil immersion objective on a CREST LFOV Spinning Disk/ C2 Confocal Microscope. Fluorescence recovery after photobleaching was achieved with a Opti-Microscan FRAP unit with 405nm laser for photobleaching. 20% laser power was used to bleach Alexa Fluor 555-labeled H1 and 15% laser power was used to bleach

Alexa Fluor 647-labeled chromatin arrays. Average H1 intensity within the bleached area was measured at 2.5 second intervals for 2 minutes. Average chromatin intensity within the bleached area was measured at 10 second intervals for 5 or 10 minutes. The signal was background subtracted and corrected for photobleaching by measuring the average intensity in an identical area of a condensate in the same field of view that had not been bleached using the following formula:

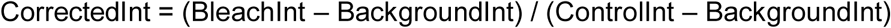

The fluorescence recovery was normalized to both the pre-bleach intensity and post-bleach intensity using the following formula:

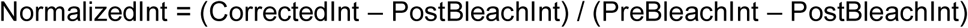

The data were then fit to the following one-phase exponential association function in GraphPad Prism to determine the rate and mobile fraction of recovery:

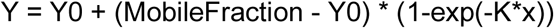

