## Supplemental Figures S1-S10 and Table S1 for "HMGB1 restores a dynamic chromatin environment in the presence of linker histone by deforming nucleosomal DNA"

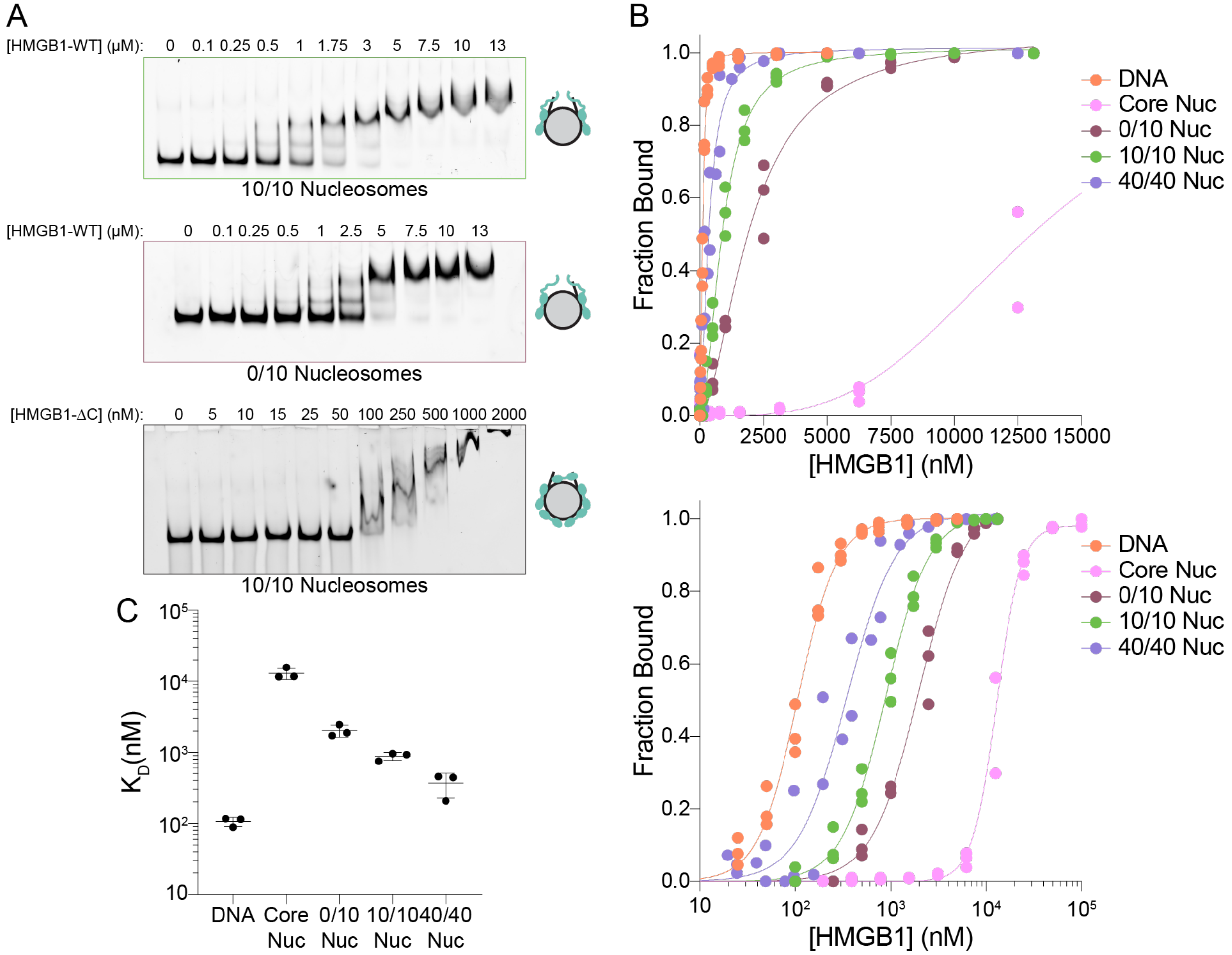


**Supplemental Figure 1. HMGB1 binding to nucleosomes and DNA**

(A) Representative images of gels depicting electrophoretic mobility shift assays (EMSA) showing binding HMGB1-WT (top 2 gels) and HMGB1-ΔC to labeled 10/10 nucleosomes (top and bottom gels) or labeled 0/10 nucleosomes (middle gel).

(B) Quantification of binding of HMGB1-WT measured by EMSA assays as shown in (A). HMGB1-WT binding to DNA (orange), core nucleosomes (pink), 0/10 nucleosomes (burgundy), 10/10 nucleosomes (green) and 40/40 nucleosomes (purple) are shown. Each condition was performed in triplicate and fit to a binding equation. Bottom panel shows the same quantification on a semi-log plot.

(C) Quantification of the $K_D$ values of HMGB1-WT binding to various DNA and nucleosome substrates. $K_D$ values are extracted from binding curve fits in (B).

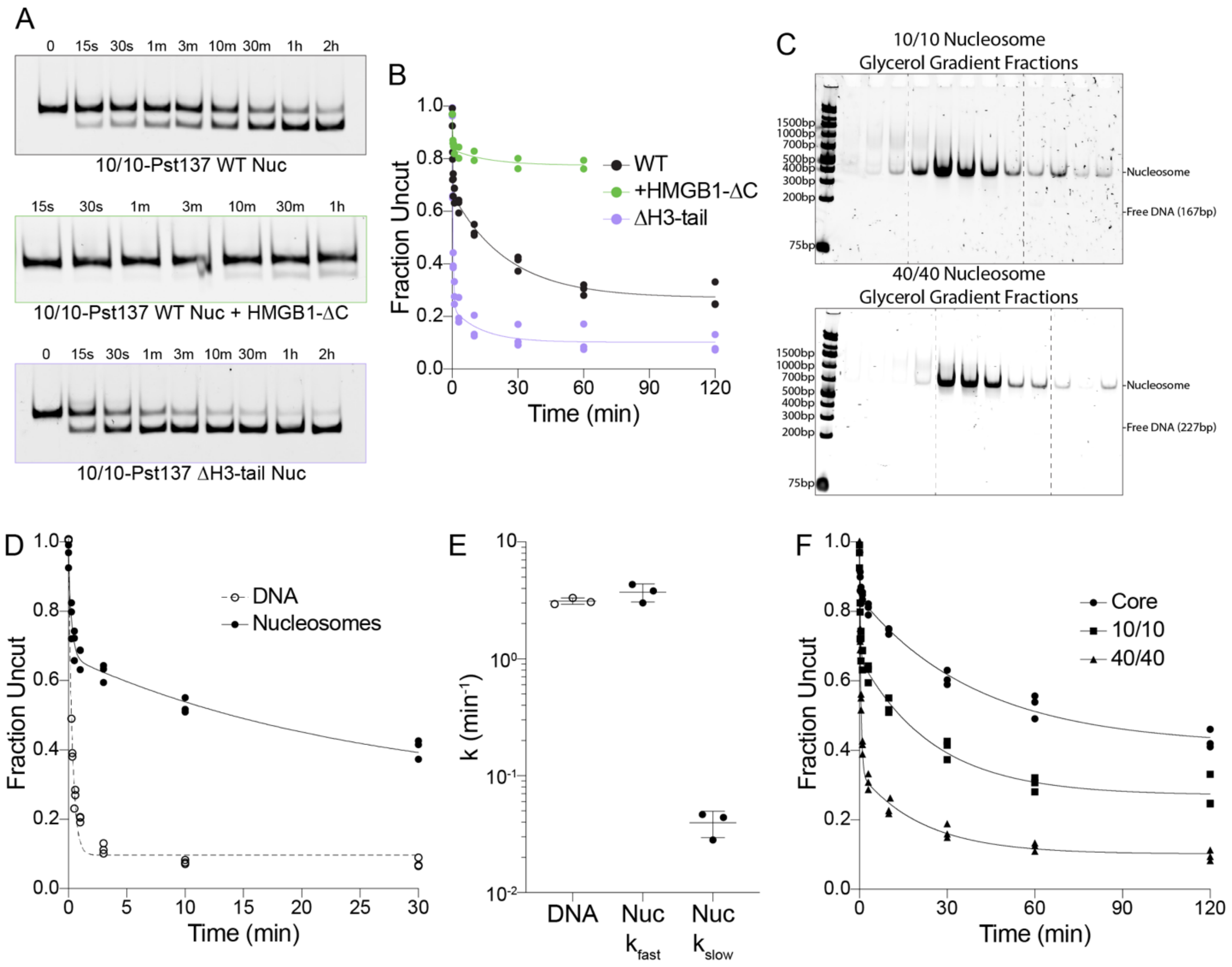


**Supplemental Figure 2. Additional REA experiments**

(A) Representative gels of REA experiments on 10/10-Pst137 WT (black) or ΔH3-tail (magenta) nucleosomes. Bottom gel is of WT nucleosomes in the presence of 100nM HMGB1-ΔC (green).
(B) Quantification of REA experiments from (A). Nucleosome alone experiments are shown in triplicate, and nucleosomes with 100nM HMGB1-ΔC in duplicate. The data are fit to a two-step exponential decay function.
(C) Representative gels of glycerol gradient fractions showing the purification of assembled 10/10 (top) and 40/40 (bottom) nucleosomes, showing the lack of free DNA. Dashed lines represent the fractions that were pooled.
(D) Quantification of REA experiments on 10/10-Pst137 WT nucleosomes (closed circles) and DNA (open circles). Nucleosome REA is fit to a two-step exponential decay function (solid line), while DNA REA is fit to a one-step exponential decay function (dashed line).
(E) Quantification of the rate constants derived from the curves in (C). Error bars represent the standard deviation of three experimental replicates.
(F) Quantification of REA experiments on Pst137 nucleosomes with either no flanking DNA (Core, circles), 10bp of flanking DNA on either side of the nucleosome (10/10, squares), or 40bp of flanking DNA on either side of the nucleosome (40/40, triangles). Each condition is fit to a two-step exponential decay function.

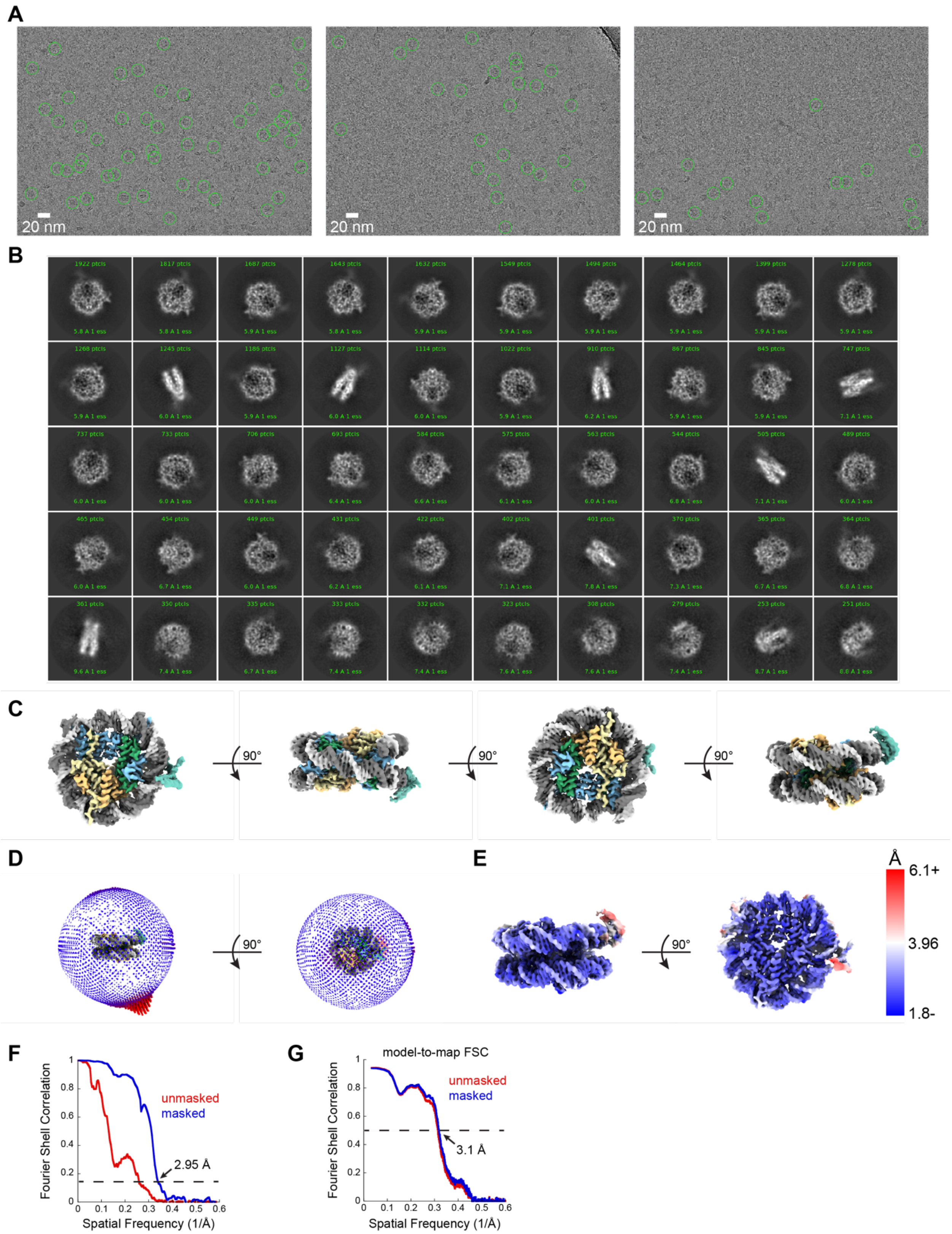


**Supplementary Figure 3. Single-particle cryo-EM of HMGB1 with 0/10 nucleosomes**

(A) Representative motion-corrected micrographs from the HMGB1 with 0/10 nucleosome dataset.

(B) Representative 2D classes of HMGB1 bound to 0/10 nucleosome.
(C) Four different views of the cryo-EM map of HMGB1 bound to 0/10 nucleosome at SHL -2 generated with non-uniform refinement in cryoSPARC v4.1.1. The structure is color-coded with histone H3 in light blue, histone H4 in green, histone H2A in yellow, histone H2B in orange, DNA strands in light/dark gray, and HMGB1 in teal.
(D) Angular distribution of particles used to generate the cryo-EM map of HMGB1 bound to 0/10 nucleosome at SHL -2 with non-uniform refinement in cryoSPARC v4.1.1.
(E) Cryo-EM map of HMGB1 bound to 0/10 nucleosome at SHL -2 generated with non-uniform refinement in cryoSPARC v4.1.1 colored by estimated local resolution determined with FSC = 0.143 cutoff in cryoSPARC v4.1.1.
(F) Unmasked (red) and masked (blue) Fourier shell correlation curves between two half-maps for the HMGB1 bound to 0/10 nucleosome at SHL -2 non-uniform refinement determined by cryoSPARC v4.1.1.
(G) Unmasked (red) and masked (blue) model-to-map Fourier shell correlation curves between the model and map for HMGB1 bound to 0/10 nucleosome at SHL -2 determined by Phenix v1.19.

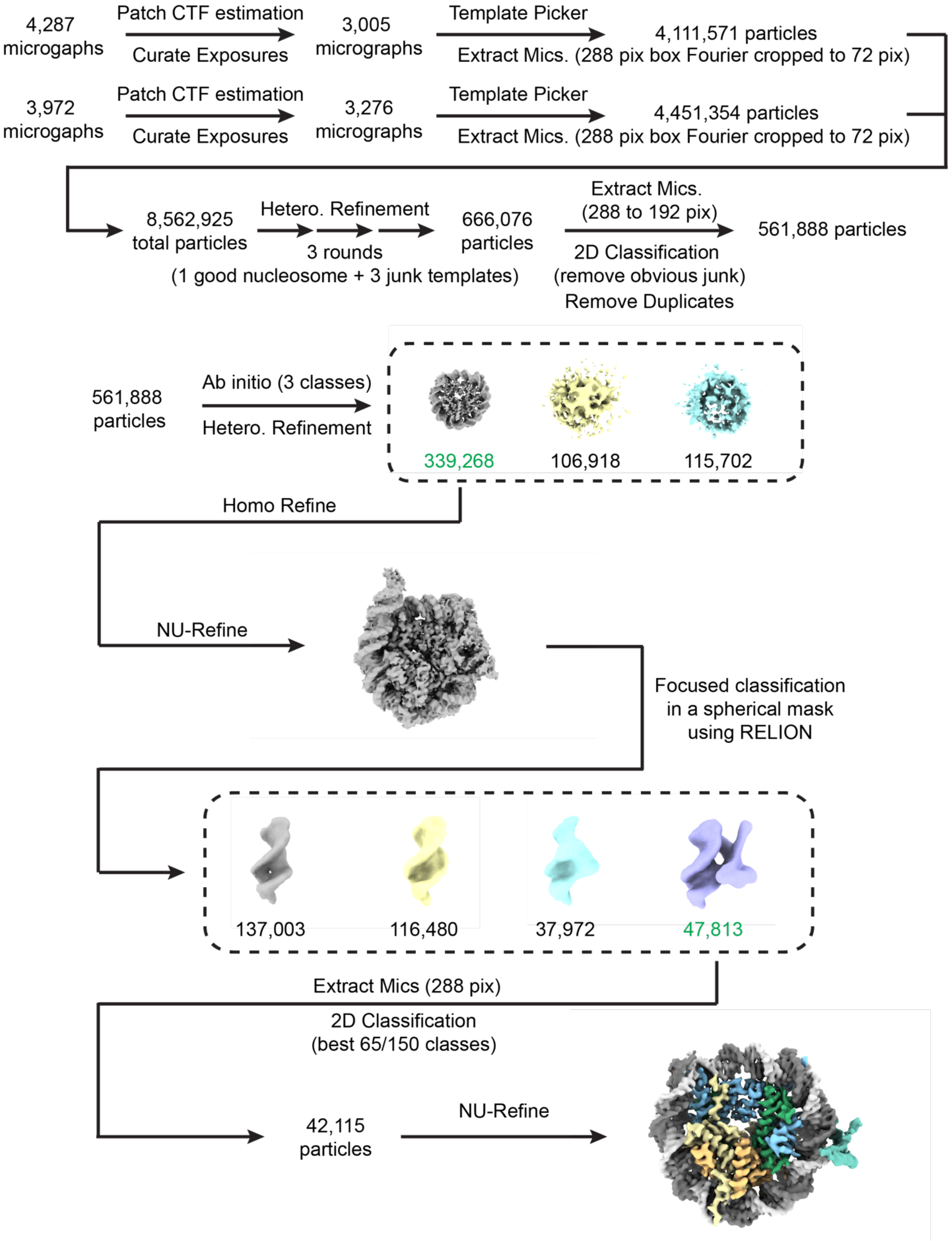


**Supplemental Figure 4. Data processing for HMGB1 with 0/10 nucleosome**

Flowchart for cryo-EM data processing of the HMGB1 with 0/10 nucleosome dataset as described in Methods. Most processing steps were performed in cryoSPARC v4.1.1 with the numbers of micrographs and particles moving into each step noted. Focused classification without alignment was performed in RELION v4.0 to improve HMGB1 density. Final non-uniform refinement was performed with cryoSPARC v4.1.1.

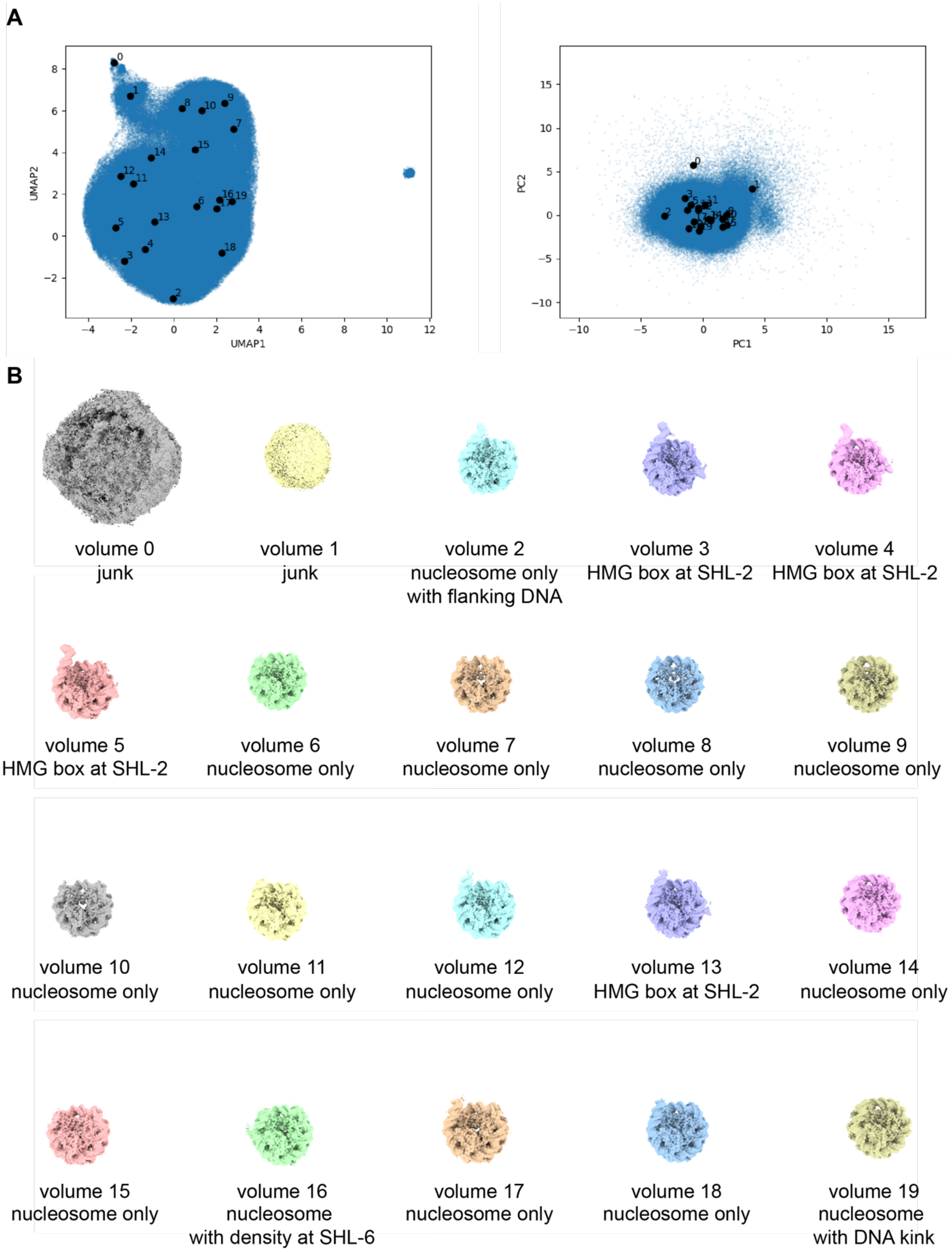


**Supplemental Figure 5. cryoDRGN analysis for HMGB1 with 0/10 nucleosome**
(A) The resulting UMAP and PCA visualizations of the particle latent embeddings after training an 8-dimensional latent variable model using cryoDRGN v1.1.2. The centers of 20 different

clusters determined by the k-means clustering algorithm to partition the latent space are annotated with a dot and corresponding cluster number.
(B) The resulting 20 density maps generated from particles partitioned by the k-means clustering algorithm. Brief descriptions of each density map are provided, with most maps corresponding to either a nucleosome-only class, nucleosome with HMGB1 bound at SHL -2, and nucleosome with kinked DNA at SHL -6 with some hint of extra density for HMGB1.

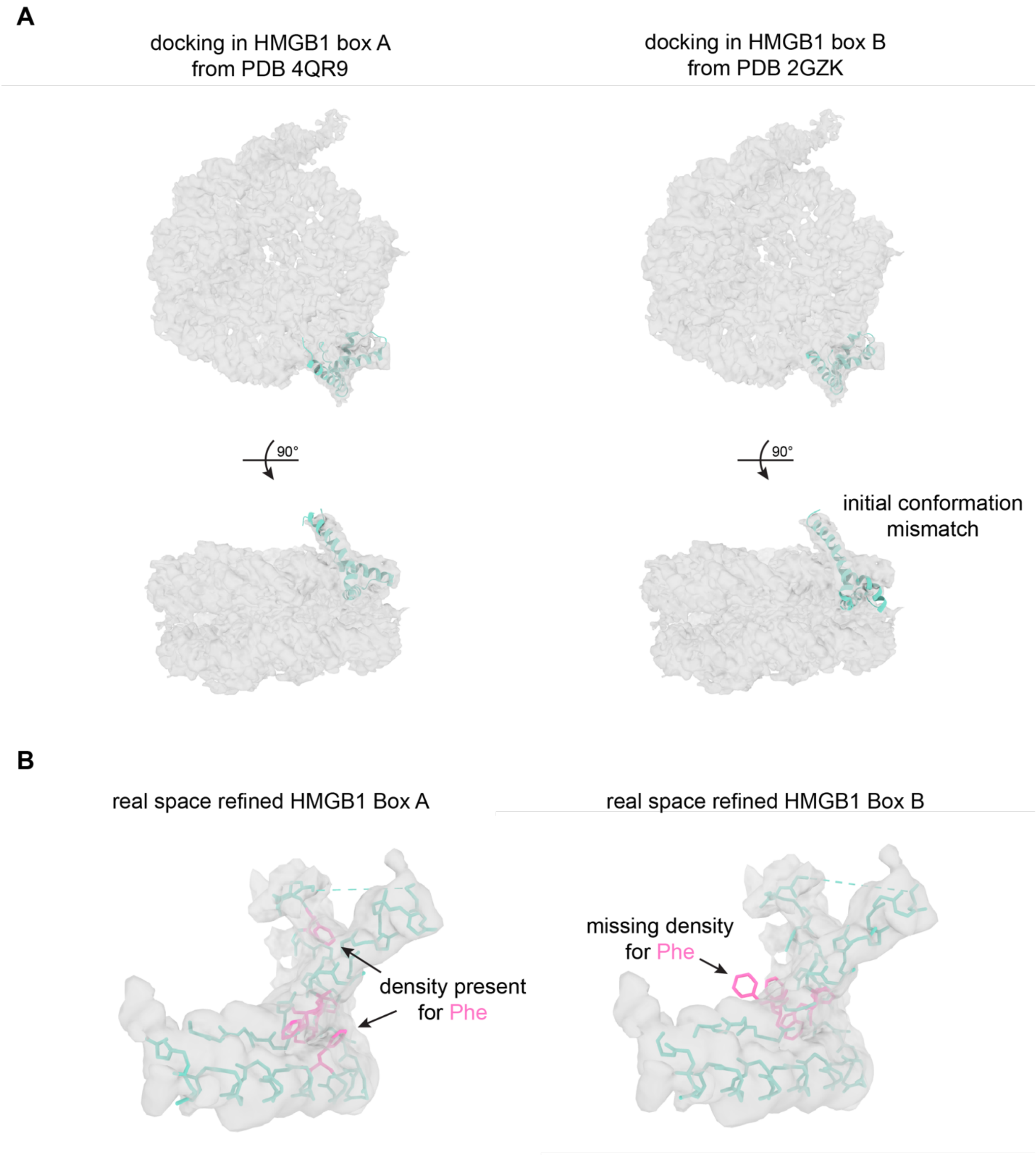


**Supplemental Figure 6. Analysis of HMGB1 density at SHL -2**
(A) The relatively low local resolution of the HMGB1 density at SHL -2 (~4 to 6 Å) precluded de novo model building in this region. Previously determined structures of HMGB1 box A bound to DNA (PDB 4QR9) and HMGB1 box B bound to DNA (PDB 2GZK) were docked into the current EM density as potential initial templates. The observed HMG box fold in our current structure more closely matches the previously determined structure of HMGB1 box A (left) and does not match the determined structure of HMGB1 box B (right).

(B) Real space refined models of HMGB1 box A (left) and HMGB1 box B (right) in the observed HMG box density at SHL -2. For simplicity, only sidechains for phenylalanine and tryptophan residues are shown in pink. Density can be observed for these bulky residues if the sequence for box A is built into the density (left) but not if the sequence for box B is built (right).

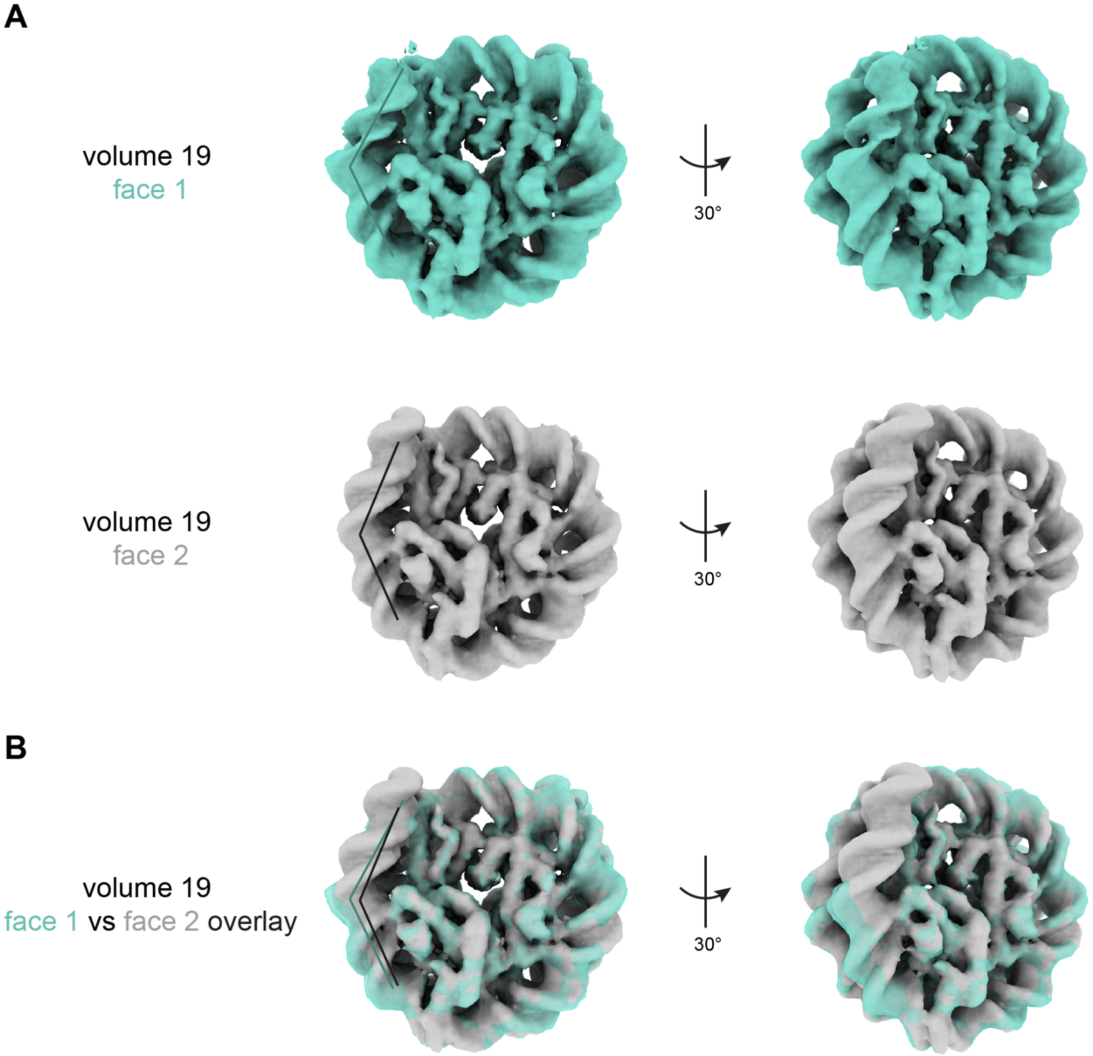


**Supplemental Figure 7. Analysis of DNA kink at SHL -6 in volume 19**
(A) Density maps of both faces of the nucleosome from volume 19 from the cryoDRGN analysis.
(B) An overlay of the density maps of both faces of the nucleosome. Face 1 (teal) shows the kink in the DNA at SHL -6.

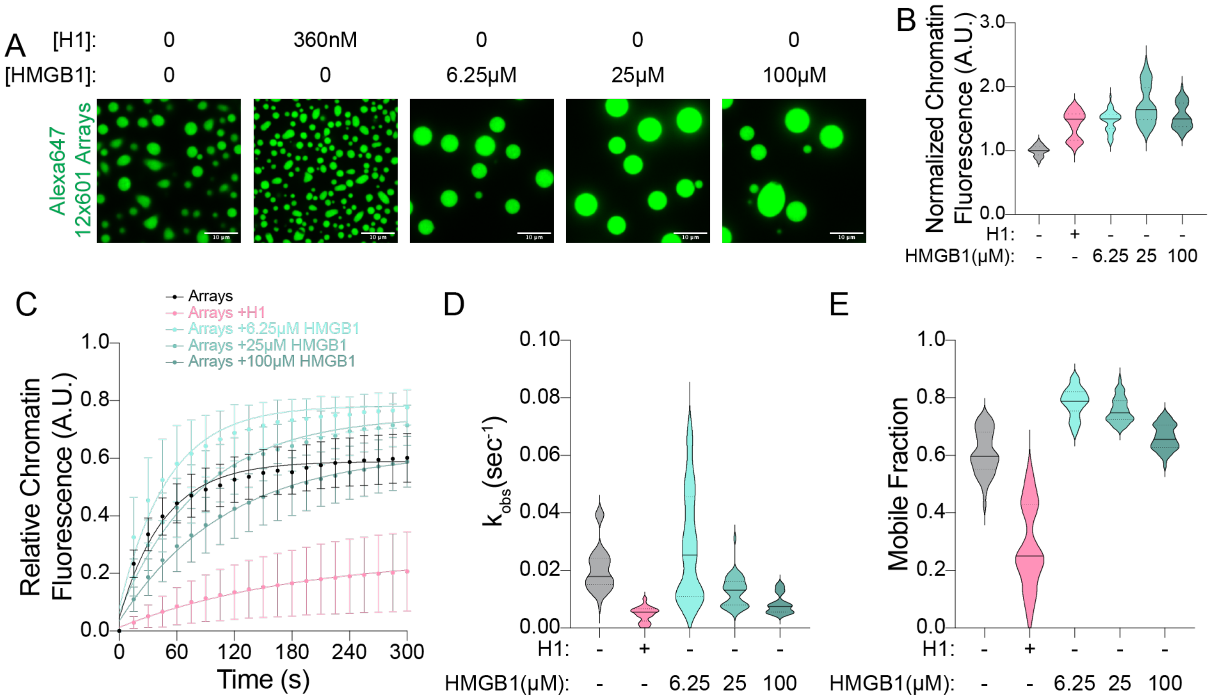


**Supplemental Figure 8. HMGB1 has minimal effect on chromatin turnover**

(A) Representative images of chromatin condensates consisting of 30nM Alexa647 12x601 chromatin arrays and various concentrations of H1 or HMGB1 as indicated. Scale bars represent 10μm.

(B) Violin plots of the quantification of the average Alexa647 12x601 chromatin arrays fluorescence intensity within chromatin condensates. All values are normalized to chromatin alone. Solid and dotted lines represent the median and interquartile values respectively. Values represent an average of three experimental replicates with n>10 condensates per replicate.

(C) Quantification of the fluorescence recovery after photobleaching of Alexa647 12x601 chromatin arrays within condensates. Values are normalized from 0 to 1 corresponding to post and pre-bleach respectively. Points represent an average of three experimental replicates with n>10 condensates per replicate. Error bars represent standard deviation of these same replicates. Data are fit to a one-phase exponentially association.

(D and E) Violin plots of the quantification of the fits to the FRAP data in (C). The rate constant for recovery ($k_{obs}$) and the plateau of recovery (Mobile Fraction) are plotted for the four different conditions in (D) and (E) respectively. Solid and dotted lines represent the median and interquartile values respectively.

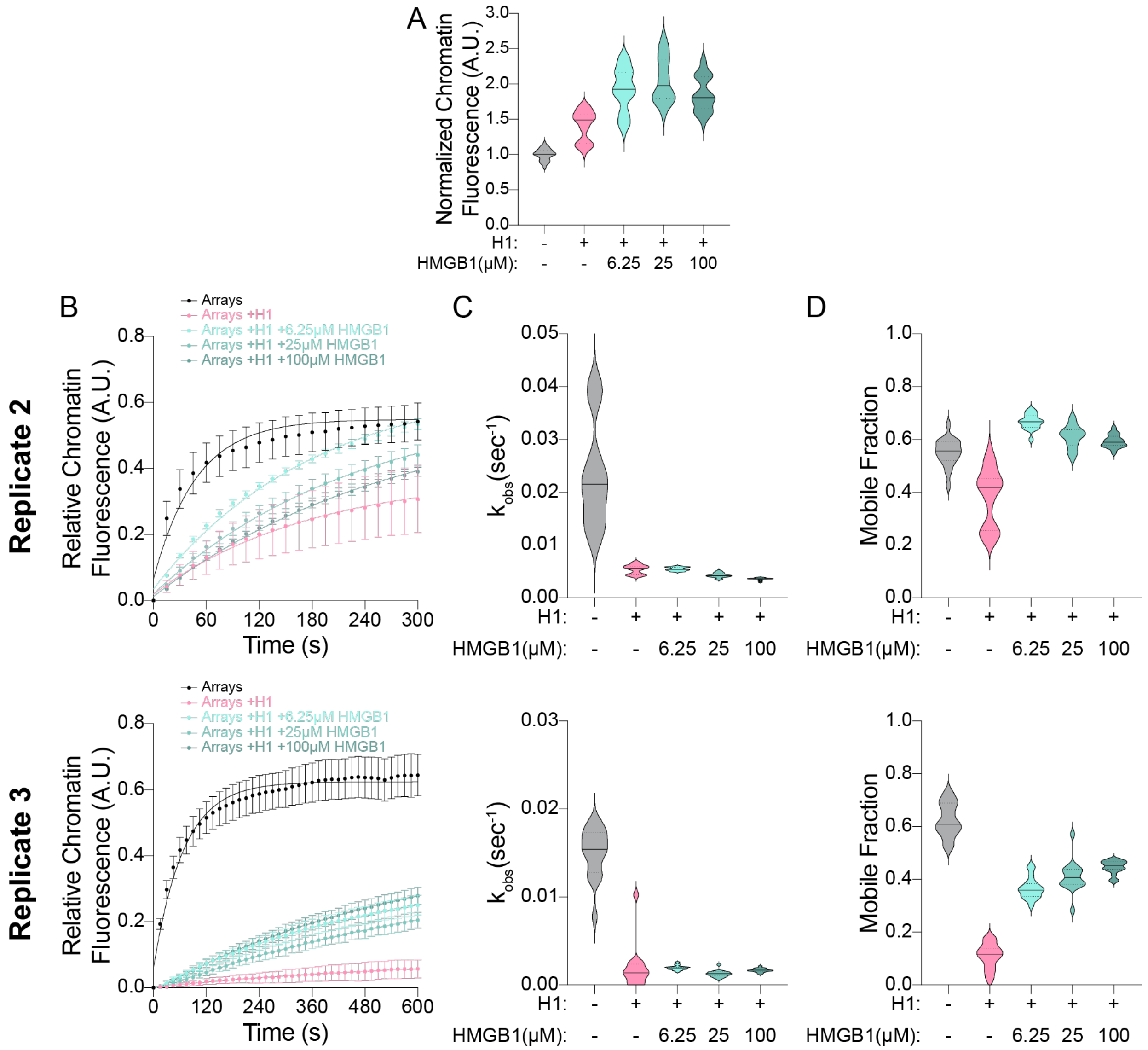


**Supplemental Figure 9. Additional replicates of H1-chromatin recovery**

(A) Violin plots of the quantification of the average Alexa647 12x601 chromatin array fluorescence intensity within chromatin condensates. All values are normalized to chromatin alone. Solid and dotted lines represent the median and interquartile values respectively. Values represent an average of three experimental replicates with n>10 condensates per replicate.

(B) Quantification of the fluorescence recovery after photobleaching of Alexa647-12x601 arrays within condensates. Values are normalized from 0 to 1 corresponding to post and pre-bleach respectively. Points represent a single experimental replicate with n>10 condensates. Error bars represent standard deviation of these same replicates. Data are fit to a one-phase exponentially association.

(C and D) Violin plots of the quantification of the fits to the FRAP data in (F). The rate constant for recovery ($k_{obs}$) and the plateau of recovery (Mobile Fraction) are plotted for the five different conditions in (G) and (H) respectively. Solid and dotted lines represent the median and interquartile values respectively.

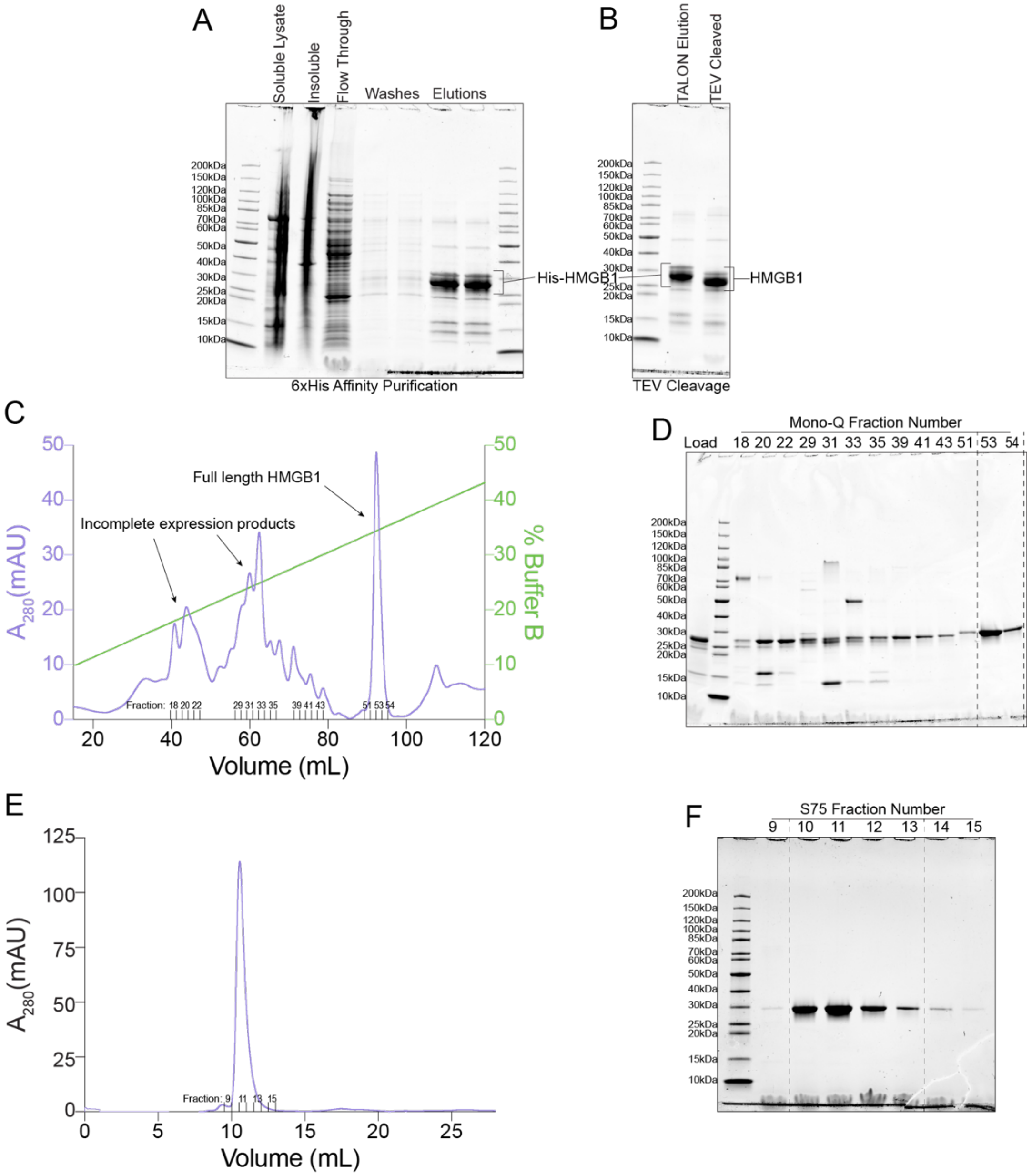


**Supplemental Figure 10. Purification of *Homo sapiens* HMGB1**

(A) A gel showing the 6xHis pulldown of His-tagged HMGB1 using TALON resin.

(B) A gel showing the cleavage of the 6xHis tag from HMGB1 using TEV protease.

(C) A chromatogram of an anion exchange MonoQ run used to purify HMGB1. $A_{280}$ is plotted on the left y-axis in purple to show where protein elutes from the column, while the % Buffer B (100% Buffer B = 1.5M KCl) is plotted on the right y-axis in green. Full length HMGB1 comes off between 32-35% Buffer B (~500mM KCl), while incomplete expression products elute at lower ionic strength, due to the lack of a complete C-terminal tail which is negatively charged. Fraction numbers that were run on a gel (D) are indicated.

(D) A gel of MonoQ fractions from (C) in addition to what was loaded onto the MonoQ column (lane 1). Dashed lines represent the fractions that were pooled.

(E) A chromatogram of a Superdex 75 Increase run used to purify HMGB1. Fractions numbers that were run on a gel (F) are indicated.

(F) A gel of Superdex 75 Increase fractions showing purified HMGB1. Dashed lines represent the fractions that were pooled.

**Supplementary Table 1: Cryo-EM data collection, model refinement and validation statistics**

| **Data collection and processing** | |
|---|---|
| Sample | HMGB1 with 0/10 nucleosome |
| Grid type | Quantifoil R1.2/1.3 200 mesh Au |
| Microscope | Titan Krios |
| Voltage | 300 kV |
| Camera | K3 |
| Energy filter | yes |
| Filter slit width (eV) | 20 |
| Magnification | 105,000 |
| Pixel size | 0.834 |
| Total electron exposure ($e^-/Å^2$) | 43 |
| Defocus range (µm) | (-0.8) - (-2.0) |
| Automation software | SerialEM |
| Micrographs after curation | 6,281 |
| Particle picker | cryoSPARC template picker |
| Total particles extracted | 8,562,925 |
| Particles in initial consensus refinement | 339,268 |
| **Reconstruction** | **EMDB XXXXX** |
| Subset | HMGB1 Box A at SHL -2 |
| Software | cryoSPARC Non-uniform Refinement |
| Final particles (dupl. removed) | 47,813 |
| Symmetry | C1 |
| Resolution, global (Å) | |
| FSC 0.5 (unmasked / masked) | 7.6 / 3.2 |
| FSC 0.143 (unmasked / masked) | 3.9 / 3.0 |
| Local resolution range (Å) | 1.8 - 6.6 |
| 3DFSC Sphericity | 0.865 |
| Sharpening B-factor ($Å^2$) | -68.8 |
| **Model Composition** | **PDB YYYY** |
| Protein residues | 824 |
| DNA | 308 |
| **Model Refinement** | |
| Refinement package | COOT/Phenix/ISOLDE |
| Model-to-map CC | 0.85 |
| R.m.s. deviations | |
| Bond lengths (Å) | 0.009 |
| Bond angles (°) | 1.459 |
| **Validation** | |
| Map-to-model FSC 0.5 | 3.1 |
| Ramachandran (%) | |
| Outliers | 0.00 |
| Allowed | 1.00 |
| Favored | 99.00 |
| MolProbity score | 0.60 |
| Poor rotamers (%) | 0.46 |
| Clashscore (all atoms) | 0.26 |
| C-beta deviations | 0.00 |
| CaBLAM outliers (%) | 0.64 |
| EMRinger score | 4.56 |
